# Single-cell resolution spatial analysis of antigen-presenting cancer-associated fibroblast niches

**DOI:** 10.1101/2024.11.15.623232

**Authors:** Xiongfeng Chen, Zhuan Zhou, Zeynep Yazgan, Luyu Xie, Francesca Rossi, Yang Liu, Bo Zhang, Patricio M. Polanco, Herbert J. Zeh, Alex C. Kim, Huocong Huang

**Author notes:** **Corresponding authors** Huocong Huang, MD, PhD Department of Surgery Department of Immunology Hamon Center for Therapeutic Oncology Research University of Texas Southwestern Medical Center 6000 Harry Hines Blvd. Dallas, TX 75390-8593, Alex C. Kim, MD, PhD Department of Surgery Hamon Center for Therapeutic Oncology Research University of Texas Southwestern Medical Center 6000 Harry Hines Blvd. Dallas, TX 75390-8593.

## Abstract

Recent studies have identified a unique subtype of cancer-associated fibroblasts (CAFs) termed antigen-presenting CAFs (apCAFs), which remain the least understood CAF subtype. To gain a comprehensive understanding of the origin and function apCAFs, we construct a fibroblast molecular atlas across 14 types of solid tumors. Our integration study unexpectedly reveals two distinct apCAF lineages present in most cancer types: one associated with mesothelial-like cells and the other with fibrocytes. Using a high-resolution single-cell spatial imaging platform, we characterize the spatial niches of these apCAF lineages. We find that mesothelial-like apCAFs are located near cancer cells, while fibrocyte-like apCAFs are associated with tertiary lymphoid structures. Additionally, we discover that both apCAF lineages can up-regulate the secreted protein SPP1, which facilitates primary tumor formation and peritoneal metastasis. Taken together, this study offers an unprecedented resolution in analyzing apCAF lineages and their spatial niches.

## Introduction

Fibroblasts are mesenchymal cells essential for preserving tissue integrity, regulating inflammatory responses and fibrosis, and facilitating wound repair (Buechler et al., 2021; Plikus et al., 2021; Younesi et al., 2024). Cancer is often described as a wound that never heals, making fibroblasts major components of the tumor microenvironment (Dvorak, 2015). In this context, they perform various functions, including the synthesis and modification of the extracellular matrix, complex signaling with cancer cells, and interactions with infiltrating immune cells (Plikus et al., 2021). Cancer-associated fibroblasts (CAFs) have thus become attractive targets for improving cancer therapies. However, the diversity and adaptability of CAFs present significant challenges in developing effective CAF-targeted treatments (Caligiuri and Tuveson, 2023; Chhabra and Weeraratna, 2023; Helms et al., 2020; Sahai et al., 2020). Single-cell RNA sequencing (scRNA-seq) has significantly advanced our understanding of fibroblast heterogeneity, revealing distinct fibroblast subtypes with specific functions in both healthy and diseased tissues (Buechler et al., 2021). Many CAF populations have been identified across different cancer types (Bartoschek et al., 2018; Elyada et al., 2019; Foster et al., 2022; Hosein et al., 2019; Hu et al., 2021; Joanito et al., 2022; Steele et al., 2020). However, the lineage relationships between CAF subtypes remain poorly understood due to the limited number of cross-tissue comparison studies and inconsistencies in CAF nomenclature (Sahai et al., 2020).

Emerging evidence suggests that phenotypically, CAFs can be classified into three major subtypes based on their molecular signatures: inflammatory CAFs (iCAFs), myofibroblastic CAFs (myCAFs), and antigen-presenting CAFs (apCAFs) (Elyada et al., 2019). This classification system provides a consistent framework to study the lineage and function of CAFs. Characterized by the expression of major histocompatibility complex class II (MHC II) molecules, apCAFs can effectively present antigens and regulate T cells (Elyada et al., 2019; Huang et al., 2022). Our previous study also found that apCAFs are derived from mesothelial progenitors in murine pancreatic ductal adenocarcinoma (PDAC) (Huang et al., 2022). However, due to their tissue- or spatial-dependent abundance, cellular plasticity, and mixed molecular signatures of immune cells and fibroblasts, apCAFs remain the least understood and characterized CAF subtype.

Major questions regarding apCAFs include their existence in human tumors, their origins in different cancer types, and the spatial niches they occupy. To address these questions, we integrated scRNA-seq data from 560 human samples, encompassing over 2.5 million cells, to construct a comprehensive fibroblast molecular atlas across 14 types of tissues and solid tumors. Our robust integration study surprisingly revealed two distinct lineages of apCAFs that exist in most tissue types, with a high abundance in specific tissues such as the peritoneum, kidney, and liver. One lineage is associated with mesothelial-like cells, as we previously reported, while the other is linked to fibrocytes (Huang et al., 2022; Schmidt et al., 2003).

To understand the stromal niches formed by the two apCAF populations, we spatially deconvolved the CAF subtype signatures derived from the atlas using a high-resolution single-cell spatial imaging platform. This approach revealed that mesothelial-like apCAFs are proximal to cancer cells, whereas fibrocyte-like apCAFs are closely associated with lymphocytes and tertiary lymphoid structures (TLS). Additionally, we discovered that apCAFs contributed to the expression of stromal SPP1, knockout of which led to a significant reduction in primary tumor weight, peritoneal metastases, and ascites formation, along with normalization of CAF formation and enhanced infiltration of lymphocytes. This work provides an analysis of apCAF lineages and niches at an unprecedented resolution, advancing our understanding of this poorly understood CAF subtype.

## Results

### CAF phenotypes are conserved across cancer types

To comprehensively characterize apCAF in human cancer, we first collected scRNA-seq datasets derived from primary human tumors and the associated normal tissues across 13 different organs including bladder (BLCA), breast (BRCA), cervix (CESC), colorectum (COLO), esophagus (ESCC), kidney (ccRCC), liver (HCC), lung (NSCLC), ovary (OV), pancreas (PDAC), prostate (PRAD), skin (BCC/SCC) and stomach (STAD) (Figure S1A and Table S1). Given that in our previous study, we found mesothelial cells could be a progenitor for apCAFs, and peritoneum is a tissue covered by a layer of mesothelial cells, we also included datasets from normal peritoneum and peritoneal metastasis (PM) (Cortes-Guiral et al., 2021; Koopmans and Rinkevich, 2018). We then performed stringent preprocessing, batch effect correction and integrative clustering with all the datasets. In the end, we obtained a high-quality scRNA-seq atlas consisting of 560 samples and over 2.5 million cells (Figure 1A and Figure S1B). We identified the major cell types across tissues by distinct signatures of each cell type: epithelial cells, T & NK cells, fibroblast-like cells, myeloid cells, endothelial cells, plasma cells, B cells and mast cells (Figure 1A and Figure S1C). By cross-comparing all the cancer types, we found that PDAC, PM and OV were most enriched with fibroblast-like cells (Figure 1B).

**Figure 1.**
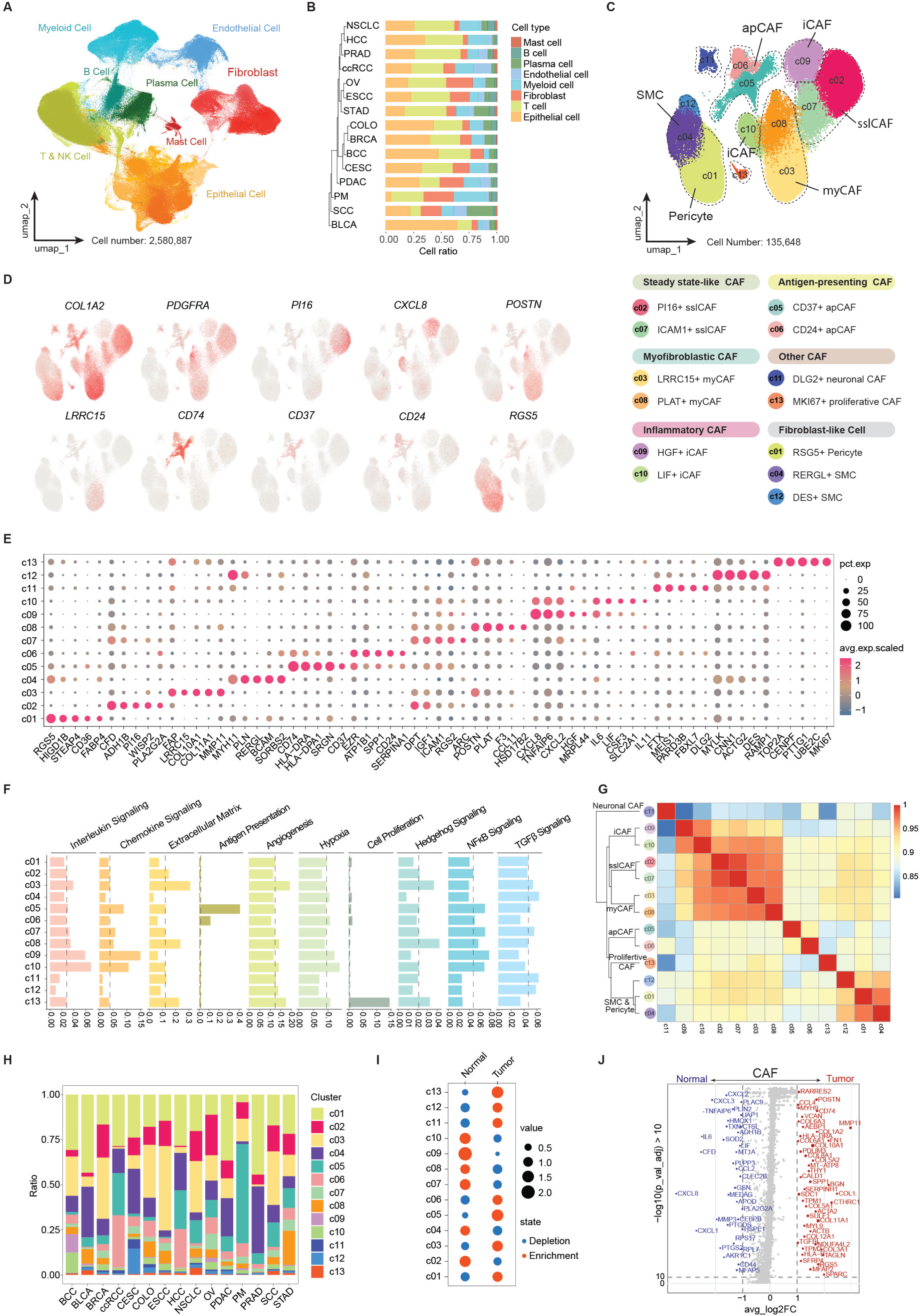
Cross-tissue comparison of CAF subtypes. (A) Integrated scRNA-seq atlas of 14 types of tissues and solid tumors. (B) Proportions of major cell types in different cancer types. (C) The 13 CAF subpopulations across tissues can be classified into steady state-like CAF (sslCAF), myofibroblastic CAF (myCAF), inflammatory CAF (iCAF), antigen-presenting CAF (apCAF), neuronal CAF, proliferative CAF and fibroblast-like cell including pericyte and SMC. (D) UMAPs of fibroblast and pericyte genes including *COL1A2* (fibroblast-associated cells), *PDGFRA* (pan-fibroblast), *PI16* (sslCAF), *CXCL8* (iCAF), *POSTN* (myCAF), *LRRC15* (myCAF), *CD74* (apCAF), *CD37* (apCAF), *CD24* (apCAF) and *RGS5* (pericyte). (E) Marker genes for each fibroblast cluster. (F) Molecular signature scores for interleukin signaling, chemokine signaling, extracellular matrix, antigen presentation, angiogenesis, hypoxia, cell proliferation, hedgehog signaling, NFκB signaling, TGFβ signaling in each CAF cluster. (G) Similarity analysis of CAF subclusters reveals apCAFs are derived from distinct lineages. (H) Proportions of CAF subclusters in different cancer types. (I) Enrichment of fibroblast subclusters in normal and tumor tissues. Tissue types with normalized values greater than 1 are categorized as enrichment, and tissue types with values less than 1 are categorized as depletion. (J) Global differentially expressed gene signatures of CAFs in normal tissues and cancer.

We then extracted the fibroblast-like cells and found they formed 13 clusters with distinct signatures, signaling and function (Figure 1C-F, Figure S1D and Table S2). Although all clusters expressed collagen gene *COL1A2*, clusters 01, 04 and 12 (c01, 04, 12) were negative for pan-fibroblast marker *PDGFRA* and *LUM* (Figure 1D and Figure S1D). Instead, c01 was marked by pericyte genes (*RGS5*, *MCAM*, *CSPG4*), while c04 and c12 were marked by smooth muscle cell (SMC) genes (*MYH11*, *PLN*) (Figure 1D-E and Figure S1D). The rest of the clusters represented all fibroblasts across tissues. We found these cells fell into 6 major subtypes based on their signatures and signaling. Among these 6 subtypes, we found iCAFs (c09, c10), myCAFs (c03, c08) and apCAFs (c05, c06). Both iCAF clusters were featured by the expression of inflammatory genes (*CXCL2*, *CXCL8*, *IL6*) and signaling pathways (interleukin, chemokine and NFkB signaling) (Figure 1D-F and Table S3). Several studies reported that hypoxia could potentiate the formation of iCAFs, and we found hypoxia signature was enriched in c09 and c10 iCAFs (Figure 1F) (Mello et al., 2022; Schworer et al., 2023). c03 and c08 expressed known myCAF genes (*POSTN*, *MMP11*) and were enriched with signaling associated with myCAFs (extracellular matrix deposition, remodeling and hedgehog signaling) (Figure 1D-F and Figure S1D). In addition, we found c03 was marked by *LRRC15*, a recently reported myCAF population contributing to immunosuppression, while c08 was marked by *PLAT* (Figure 1D-E) (Krishnamurty et al., 2022). In contrast, we found commonly used myCAF activation markers such as *ACTA2* and *FAP* were not specific (Figure S1D). Importantly, we found two robust clusters of apCAFs (c05 and c06) featured by MHC II gene (*CD74*, *HLA-DRA*, *HLA-DPA1*) expression and antigen presentation function (Figure 1D-F). Each apCAF population also had differentially expressed genes. For example, c05 expressed *SRGN*, *CD37* and complement C1q genes (*C1QA*, *C1QB*, *C1QC*) while c06 expressed *CD24* and *SERPINA1* (Figure 1D-E and Figure S1D). In addition to iCAFs, myCAFs and apCAFs, we found c02 and c07 expressed the universal resident fibroblast marker *DPT* and *CD34*, with c02 also expressing the progenitor marker *PI16*, suggesting these two clusters were the normal tissue resident fibroblasts, serving as possible progenitors for iCAFs and myCAFs (Figure 1D-E and Figure S1D) (Buechler et al., 2021). In a recent study that cross-compares PDAC and breast cancer, these fibroblasts are reported as steady-state-like CAFs (sslCAFs) (Foster et al., 2022). c11 and c13 were the minor CAF populations, carrying a distinct neuronal (*DGL2*, *PIEZO2*) and proliferative (*MKI67*) signature respectively (Figure 1E and Figure S1D). Among all the CAF populations, we found c03 myCAFs, c07 sslCAFs, c11 neuronal CAFs and SMCs were most active for TGFβ signaling (Figure 1F).

Phylogenetic analysis showed that sslCAFs, iCAFs and myCAFs belonged to the same branch while the two apCAF populations formed a distinct branch, suggesting different progenitors for the apCAF lineages (Figure 1G). We then analyzed the distribution of fibroblast subtypes across tissues. Surprisingly, we found that although there were proportional differences, the constituents of CAF subtypes were conserved regardless of cancer types (Figure 1H). One exception was c09 iCAFs, which was marked by a higher level of *HGF* (Figure 1E). We found this iCAF population was mostly enriched in skin and ovarian cancer. We further investigated the relative percentage alteration of each fibroblast cluster between normal tissues and tumors (Figure 1I). We found that *LRRC15^+^*myCAFs (c03), both apCAF subtypes (c05, c06), neuronal CAFs and proliferative CAFs were the only CAF subtypes that increased in tumors. In contrast, both steady-state like (c02, c07) and inflammatory fibroblasts (c09, c10) relatively decreased in tumors. These data suggest that steady-state like and inflammatory fibroblasts are the major fibroblast populations in normal tissues and they probably contribute to the formation of myCAFs during tumor progression. Consistently, we also found that there was an increase of myofibroblastic and antigen-presenting signatures and a decrease of inflammatory signature in fibroblasts globally after tumors form (Figure 1J).

### Identification of two distinct lineages of apCAFs

To further interrogate the origin and molecular features of the two apCAF populations, we extracted c05 and c06 from the atlas and performed re-clustering of the cells, which revealed two distinct lineages consisting of 4 subclusters (Figure 2A and Figure S1E). One lineage (clusters 0, 3) carried a mesothelial-like signature marked by mesothelial genes (*MSLN, UPK3B, KRT19*) (Figure 2B) (Huang et al., 2022). Moreover, this lineage also expressed *CD24*, a recently reported innate immune checkpoint and involved in the modulation of cell stemness and differentiation (Figure S1F) (Barkal et al., 2019). These data are consistent with our previous study demonstrating the mesothelial origin of apCAFs. Moreover, we discovered apCAFs could be derived from another lineage of cells (clusters 1, 2) that carried fibrocyte features (Figure 2A). Fibrocytes have been reported to be derived from a hematopoietic origin and expressed hematopoietic markers such as CD45 and fibroblastic markers such as extracellular matrix proteins at the same time (Direkze et al., 2004; Han et al., 2023; Phillips et al., 2004; Schmidt et al., 2003). They are known to be able to migrate to wound sites and facilitate inflammation, fibrotic response and wound healing. We found the fibrocyte-like lineage of apCAFs were marked by many hematopoietic genes such as *PTPRC* (gene for CD45), *CD37* and *CD52* (Figure 2B and Figure S1F). Therefore, in this study, we named the mesothelial-associated lineage M-apCAFs and the fibrocyte-associated lineage F-apCAFs. We then performed pseudotime analysis, which suggested differentiation occurred in the two progenitors of apCAFs (Figure 2C). We found that M-apCAFs down-regulated the mesothelial gene *MSLN*, F-apCAFs down-regulated *PTPRC* while both lineages up-regulated MHC II genes (*CD74*, *HLA-DRA*) during the differentiation. Pathway analysis of F-apCAFs (Figure 2D) and M-apCAFs (Figure 2E) suggested that both apCAF lineages were heavily involved in immune-regulating activities.

**Figure 2.**
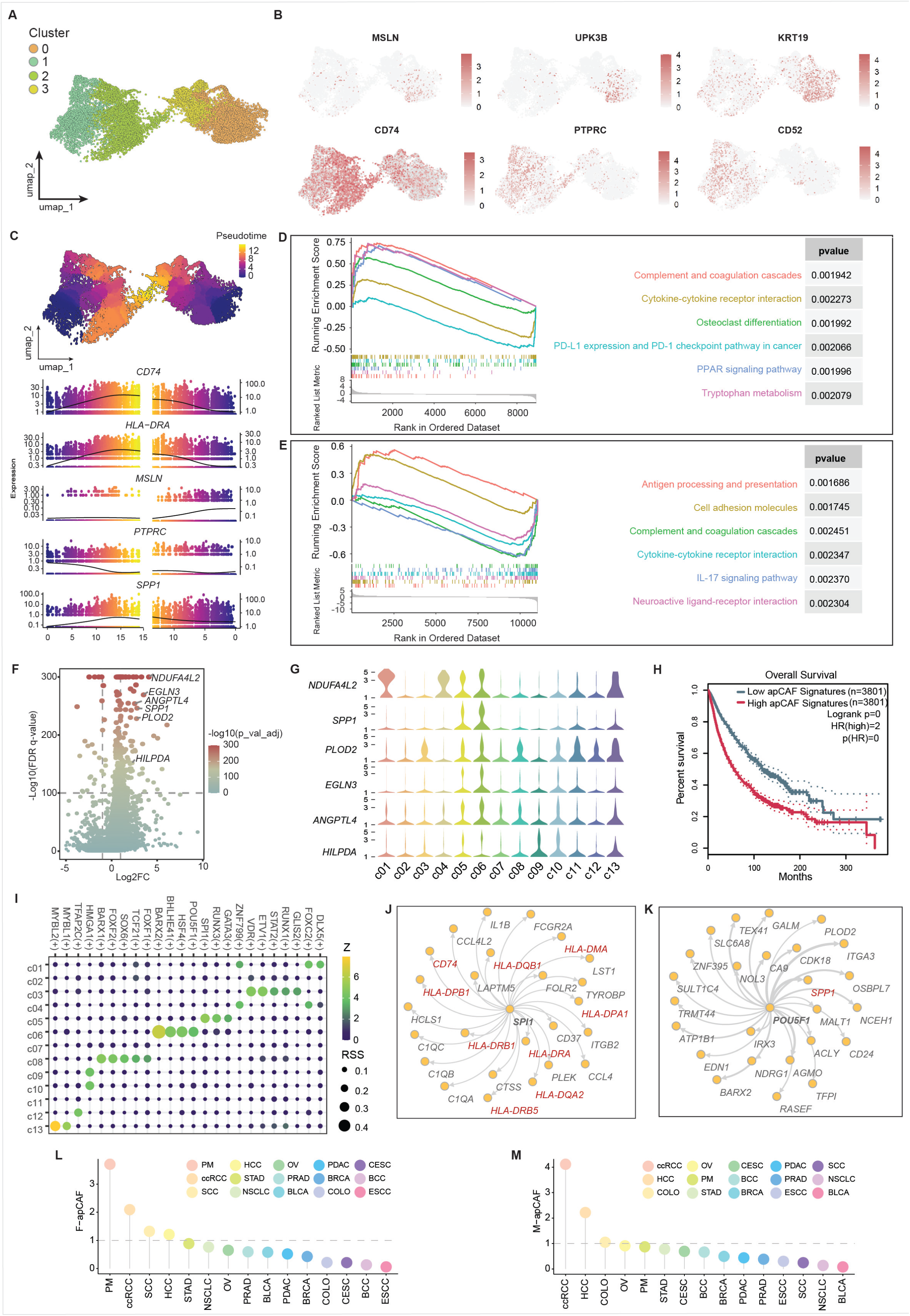
Two distinct lineages of apCAFs. (A) All apCAFs marked by MHC II molecule expression are extracted from Figure 1C and re-clustered, revealing four apCAF subclusters. (B) UMAPs of signature genes of apCAF subclusters including *CD74*, *MSLN*, *UPK3B*, *KRT19*, *PTPRC*, *CD52* (C) Pseudotime analysis reveals two distinct trajectories of apCAFs. Expression of *CD74*, *HLA-DRA*, *MSLN*, *PTPRC* and *SPP1* along the trajectories are shown. (D) Up-regulated genes in the F-apCAF lineage (subcluster 2 vs 1) are used to perform GSEA pathway analysis. Significant pathways are shown. (E) Up-regulated genes in the M-apCAF lineage (subcluster 3 vs 0) are used to perform GSEA pathway analysis. Significant pathways are shown. (F) Differentially expressed genes in apCAFs in cancer compared to normal tissues. Six most up-regulated and robustly expressed genes are identified: *NDUFA4L2*, *SPP1*, *PLOD2*, *EGLN3*, *ANGPTL4*, *HILPDA*. (G) Expression of *NDUFA4L2*, *SPP1*, *PLOD2*, *EGLN3*, *ANGPTL4*, *HILPDA* in each CAF subcluster. (H) Combined overall survival of the 14 types of cancer with the six-gene signature (*NDUFA4L2*, *SPP1*, *PLOD2*, *EGLN3*, *ANGPTL4*, *HILPDA*). (I) Regulatory network of transcription factors in each CAF subcluster revealed by SCENIC algorithm. (J) Regulatory network of genes by *SPI1* in F-apCAFs. (K) Regulatory network of genes by *POU5F1* in M-apCAFs. (L) Abundance of F-apCAFs in different cancer types. (M) Abundance of M-apCAFs in different cancer types.

We compared the differentially expressed genes between normal and cancer tissues in the two apCAF lineages and identified five most up-regulated and robustly expressed genes when tumors formed (*NDUFA4L2*, *SPP1*, *PLOD2*, *EGLN3*, *ANGPTL4*, *HILPDA*) (Figure 2F-G). This six-gene signature significantly correlated with a poor survival in cancer patients (Figure 2H). We inquired the CAF atlas about the expression of these six genes and found that F-apCAFs (c05) and M-apCAFs (c06) were the major source of *SPP1* among all CAF populations (Figure 2G). Additionally, *SPP1* was up-regulated during the differentiation of both apCAF lineages (Figure 2C). The expression of *SPP1* itself also served as a bad prognostic factor for patient survival (Figure S1G).

To understand the regulatory network in apCAFs, we performed SCENIC analysis which revealed distinct transcription factors that drove the gene program for F-apCAFs (*SPI1*, *RUNX3*, *GATA3*) and M-apCAFs (*BARX2*, *BHLHE41*, *HSF4*, *POU5F1*) (Figure 2I and Figure S1H-I) (Aibar et al., 2017). These transcription factors might directly regulate the transcription of the key apCAF signatures. For example, we found *SPI1* could directly promote the expression of MHC II genes (*CD74*, *HLA-DRA*, *HLA-DRB1*, *HLA-DRB5*, *HLA-DPA1*, *HLA-DPB1*, *HLA-DQA2*, *HLA-DQB1*, *HLA-DMA*) in F-apCAFs (Figure 2J). We also found *POU5F1* (gene for OCT4, the major transcription factor regulates stemness) transcribed *SPP1* in M-apCAFs (Figure 2K) (Takahashi and Yamanaka, 2006). Moreover, ChIP-seq of OCT4 showed that it directly bound to the promoter region of *SPP1* (Figure S1J). These data suggest that the expression of *SPP1* might be regulated by a regenerative program induced by the wound signal in the tumor microenvironment. We then analyzed the abundance of the two apCAF lineages across cancer types, and found F-apCAFs were most enriched in PM, ccRCC, SCC, HCC and STAD, while M-apCAFs were most enriched in ccRCC, HCC, COLO, OV and PM (Figure 2L-M).

### Stromal heterogeneity of peritoneal metastasis

Peritoneum is a complex tissue that consists of numerous stromal cells, including mesothelial cells, fibroblasts, endothelial cells and immune cells (Cortes-Guiral et al., 2021). This specific microenvironment may facilitate the formation of apCAFs during metastasis. Since PM is a manifestation that was enriched with apCAFs, we attempted to use this model to further understand apCAFs. One of the most common origins of PM is colon cancer, which occurs in about 20% of colon cancer patients. Recent scRNA-seq study has reported that cancer cells from primary colon cancer can be classified into two intrinsic epithelial phenotypes (iCMS2 and iCMS3) based on common epithelial developmental programs (Joanito et al., 2022). Notably, iCMS3 is characterized by the expression of goblet cell and mucin genes and mostly overlaps with a mucinous tumor pathological diagnosis. Given the cancer cell heterogeneity in primary tumors, we first tried to understand whether such heterogeneity persisted in the setting of PM and the associated CAF signatures using an unbiased whole-genome spatial transcriptomics approach.

We collected tissue specimens from 8 patients with PM originating from primary colon cancer. Employing the Nanostring GeoMx human whole transcriptome atlas spatial RNA profiling platform, we characterized the molecular signatures of tumor epithelium and adjacent CAF in PM (Figure 3A-B). Spatial RNA signatures of the epithelial regions of interest (ROIs) underwent deconvolution and principal component analysis, revealing two distinct molecular subtypes of epithelium carrying iCMS2 or iCMS3 signature (Figure 3C-D). Cell-type specific enrichment analysis revealed that iCMS2 epithelial cells closely resembled enterocytes, whereas iCMS3 epithelial cells resembled goblet cells, indicating the presence of two distinct cell lineage programs within PM cancer cells (Figure 3E). To ascertain the predictive and prognostic significance of these epithelial subtypes, our focus shifted to primary colon cancer cases at an advanced T4 stage, where the tumor had breached the outer lining of the colon and initiated spread into the peritoneum or neighboring organs (Cortes-Guiral et al., 2021). In this context, we conducted immunohistochemical (IHC) staining on a cohort of T4 samples using the iCMS3 marker SPINK4 (Figure 3F and Figure S2A-B). Our findings revealed a significant association between the presence of SPINK4^+^ invasion fronts and an elevated risk of PM occurrence in T4 tumors compared to SPINK4^-^ counterparts (relative risk=16.5, P=0.005) (Figure 3G). This underscores the significance of iCMS3 epithelial cells as potential drivers of PM.

**Figure 3.**
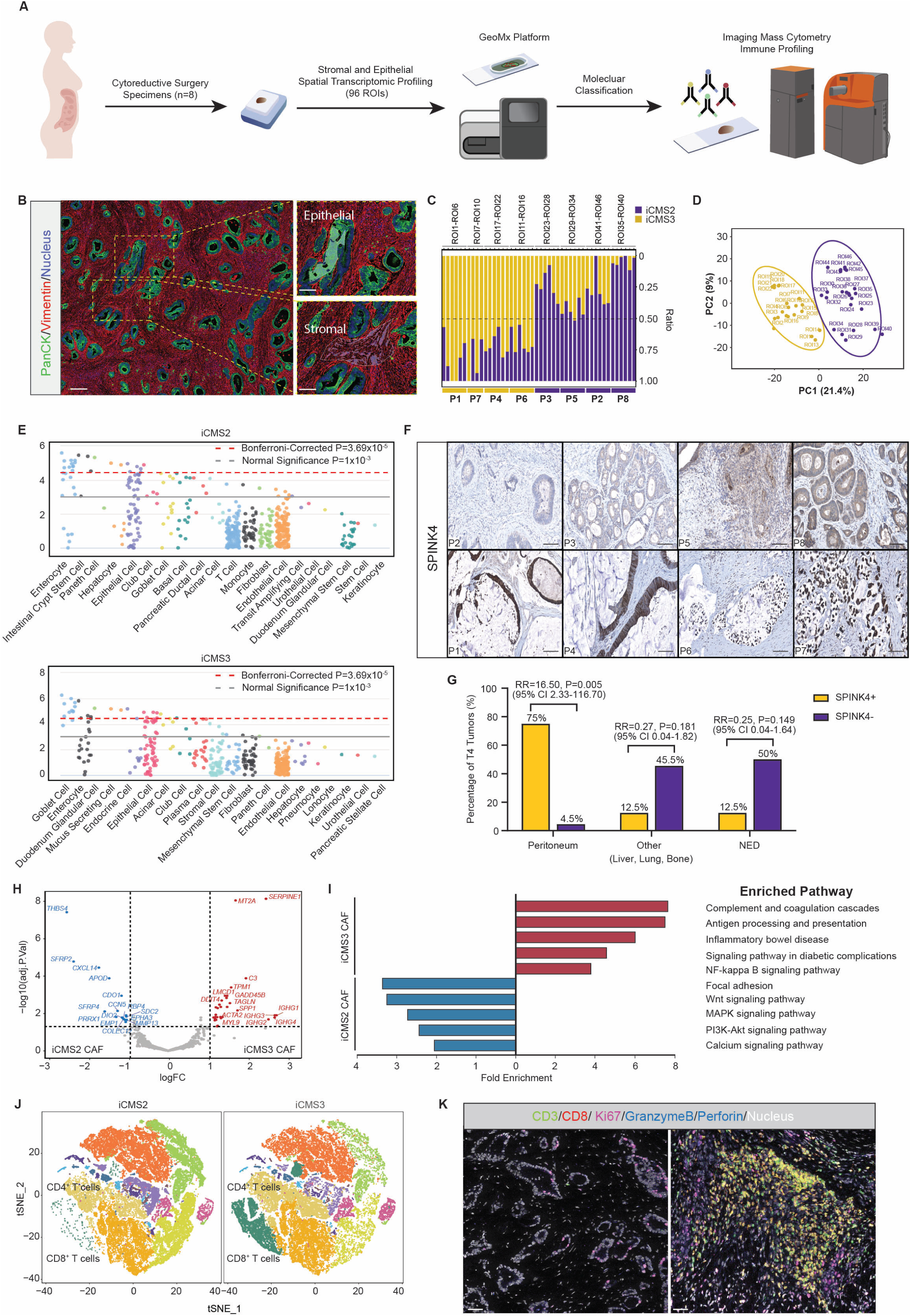
Molecular classification of PM stroma. (A) Scheme of PM molecular classification by GeoMx spatial transcriptomics and imaging mass cytometry. (B) Representative image of morphology maker staining for pan-cytokeratin (PanCK), vimentin and nuclei from the Nanostring GeoMx assay. (C) Deconvolution analysis of GeoMx spatial RNA data from PM with iCMS2 and iCMS3 gene signatures. Epithelial ROIs among the 8 patient samples can be classified into iCMS2-like (patient 2, 3, 5, 8 (P2, P3, P5, P8)) or iCMS3-like (patient 1, 4, 6, 7 (P1, P4, P6, P7)). (D) PCA analysis of all ROIs of the iCMS2 and iCMS3 groups of PM. (E) Web-based cell-type specific enrichment analysis with differentially expressing genes of iCMS2 and iCMS3 PM. (F) IHC staining for SPINK4 in iCMS2 (patient 2, 3, 5, 8 (P2, P3, P5, P8)) and iCMS3 (patient 1, 4, 6, 7 (P1, P4, P6, P7)) PM groups. Scale bars, 50 μm. (G) Percentages and relative risks of PM or metastases in other sites (liver, lung, bone) in T4 colon cancer that carry SPINK4^+^ or SPINK4^-^ invasion borders. RR, relative risk, CI, confidence interval, NED, no evidence of disease. (H) Volcano plot analysis showing top differentially expressed genes of the iCMS2 and iCMS3 CAFs in PM. (I) Pathway analysis of differentially expressed genes from iCMS2 or iCMS3 CAF in PM. (J) Imaging mass cytometry is performed in iCMS2 and iCMS3 PM samples (n=3/group, 2 images/sample) on the Hyperion platform with a total of 20 immune markers. Imaging information from all 6 samples is integrated and tSNE analysis is conducted and split into two tSNE plots by iCMS subtypes. (K) Representative images of iCMS2 and iCMS3 PM samples from imaging mass cytometry assay including T cell markers (CD3, CD8, granzyme B, perforin and Ki67).

Next, we analyzed the transcriptomic information of tumor cell-adjacent CAFs. When we grouped CAFs into iCMS2 and iCMS3 categories, we found iCMS3 CAFs displayed enrichment in antigen presentation and inflammation-related pathways, indicating iCMS3 PM might be enriched with apCAFs (Figure 3H-I). Considering the unique inflammatory profiles observed in iCMS3 PM, we speculated potential differences in immune cell infiltration between iCMS2 PM and iCMS3 PM. To explore this, we employed imaging mass cytometry in both subtypes. Our analysis unveiled a significant rise in T cells within iCMS3 PM, however, with a lack of key activation markers like granzyme B, perforin, IFNγ, and Ki67 (Figure 3J-K and Figure S2C-D). This indicates the presence of an immune-inhibitory signal within the inflammatory milieu, impeding effective T cell activation despite their abundance. We validated these results with IHC staining for T cells in a larger cohort of PM samples (Figure S2E-F).

### Spatial analysis of F-apCAF niches

To further dissect the spatial distribution of apCAFs and the associated niches at a single cell resolution, we exploited the Xenium In Situ high-plex spatial imaging platform. We designed a fully customized panel of probes consisting of 480 genes based on the scRNA-seq atlas we generated (Figure 1 and Table S4). We then performed the Xenium assays with 8 human PM samples of colon origin including one PM-adjacent tissue. We obtained a total of 799,455 high quality cells across samples. Unsupervised clustering analysis resulted in the identification of major cell types including CAFs, SMCs, endothelial cells, lymphocytes, myeloid cells, iCMS2 and iCMS3 cancer cells (Figure 4A-B). Among the 8 PM samples, 3 samples carried a distinct iCMS3 epithelial signature (PM1, PM2, PM3), 2 carried iCMS2 signature (PM6, PM7), while the other 2 samples had a mix of both iCMS2 and iCMS3 signatures (PM4, PM5) (Figure S3A). We found that the presence of iCMS3 cancer cells was associated with an increase infiltration of lymphocytes and the formation of tertiary lymphoid structures (TLS) (Figure 4C and Figure S3A), consistent with the stromal signatures revealed by the GeoMx assay (Figure 3).

**Figure 4.**
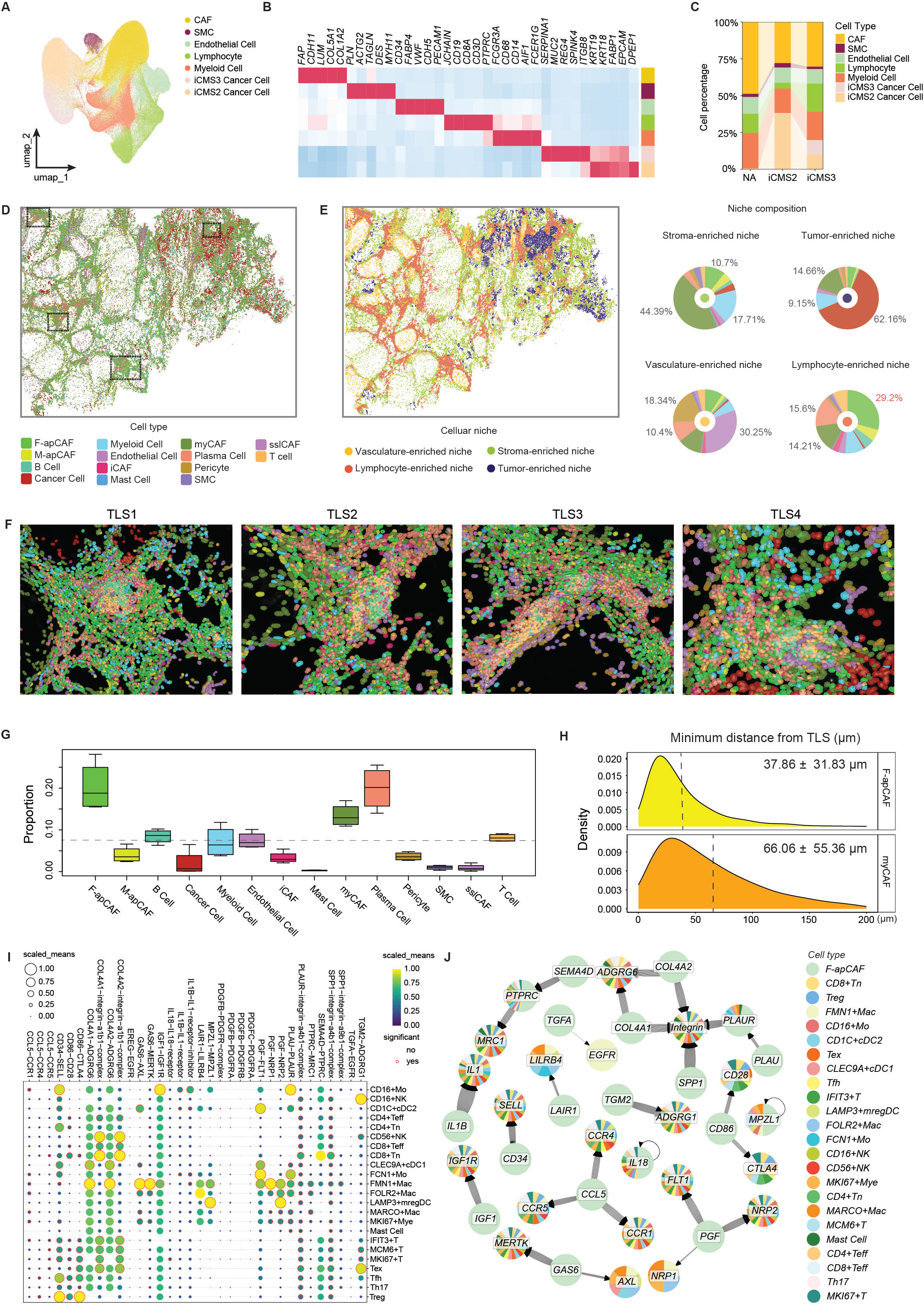
Spatial analysis of F-apCAF niches. (A) UMAP of major cell types across the 8 human PM samples identified from the Xenium assays. (B) Marker genes of the major cell types in PM. (C) Proportions of the major cell types in the non-tumor adjacent (NA), iCMS2 and iCMS3 PM samples. (D) Robust cell type decomposition is performed in iCMS3 PM to deconvolve the Xenium data into cell types using our pan-cancer scRNA-seq atlas as reference. (E) Spatial niches are computed by calculating the cell type composition of each cell using the k-nearest neighbor algorithm. Cells with similar neighborhoods are grouped into spatial niches using k-means clustering. Percentages of cell types within each niche are shown. (F) Visualization of cellular components in four TLS regions we use for quantification. (G)Quantification of the proportions of cell types present in the TLS regions. (H) Cell density and spatial distance analysis of F-apCAFs or myCAFs from TLS is conducted in the TLS regions. (I) Ligand-receptor interaction analysis between F-apCAFs (ligands) and different populations of immune cells (receptors). Lymphocyte and myeloid cell subtypes are identified from the pan-cancer scRNA-seq atlas. (J) Significant ligand-receptor pairs are shown between F-apCAF-derived ligands and receptors on immune cells.

Next, we conducted the robust cell type decomposition to deconvolve spatial transcriptomic data into cell types using the reference constructed from our pan-cancer scRNA-seq atlas (Figure 4D) and then cellular niche composition analysis (Figure 4E). We found that in the iCMS3 PM samples with marked TLS formation, four niches could be identified (vasculature-, lymphocyte-, stroma- and tumor-enriched niches). Niche composition showed that myCAFs and myeloid cells were the major stromal cells in the stroma- and tumor-enriched niches, while vasculature-enriched niches mostly consisted of endothelial cells and pericytes. Remarkably, we found that F-apCAFs were the most abundant cell type in the lymphocyte-enriched niches (29.2%) (Figure 4E). We then selected four TLS areas and further quantified the proportions of each cell type (Figure 4F). We found across the TLS areas, F-apCAFs were the most prevalent CAF subtype (Figure 4G). Although myCAFs also contributed to a major cell proportion in the TLS, we found F-apCAFs had higher density and closer interaction with TLS compared to myCAFs (average distance 37.86 µm vs 66.06 µm) (Figure 4H). To investigate the interaction between F-apCAFs and immune cells, we first identified the myeloid cell and lymphocyte subtypes from our pan-cancer scRNA-seq atlas based on and marker genes for each cluster and published literatures (Figure S3B-E) (Andreatta et al., 2021; Cheng et al., 2021; Chu et al., 2023; Tang et al., 2023). We then performed ligand-receptor interaction analysis with F-apCAF-derived ligands and immune cell-expressing receptors. The results revealed many important ligands that mediated the immune-regulating function of F-apCAFs (Figure 4I-J). For example, CCL5, which is a chemoattractant for T cells and monocytes, probably contributes to the formation of TLS (Huffman et al., 2020). However, immunosuppressive ligands were also found such as *SPP1* and *LAIR1*, suggesting the complex role of F-apCAFs in inflamed tumors (Klement et al., 2018; Zhang, 2022).

### Spatial analysis of M-apCAF niches

Compared to F-apCAFs, the spatial distribution of M-apCAFs was more context-dependent (Figure S4). Peritoneum is lined by a layer of mesothelial cells, therefore, when cancer cells metastasize onto peritoneum, the direct interaction with mesothelial cells may induce the formation of M-apCAFs (Cortes-Guiral et al., 2021; Huang et al., 2022). M-apCAFs express cytokeratin genes (Figure 2B). In fact, when we performed IHC staining for cytokeratin, we could identify robust infiltration of cytokeratin^+^ CAFs into the PM stroma which tended to locate near the cytokeratin^+^ mesothelium (Figure S5A). Further multiplex IHC staining with cancer cell marker (SPINK4) ruled out these CAFs were cancer cell-derived (Figure S5B). In the PM samples with marked cytokeratin^+^ CAF infiltration, when we conducted cell type deconvolution and niche composition analysis with the spatial transcriptomic data, we identified distinct mesothelial-enriched niches in addition to stroma-, vasculature- and tumor-enriched niches (Figure 5A-B). Within these mesothelial-enriched niches, M-apCAFs (23.89%), myeloid cells (15.48%) and T cells (12.38%) were the major stromal cells (Figure 5B). We also found some mesothelial-enriched niches were continuation of adjacent normal mesothelium (marked by normal mesothelial genes *MSLN*, *UPK3B*) (Figure S5C), suggesting the mesothelial cell to M-apCAF differentiation. Moreover, most of the mesothelial-enriched niches were near tumor epithelium (Figure 5C). These data suggest M-apCAFs may be involved in close interaction with immune cells and cancer cells.

**Figure 5.**
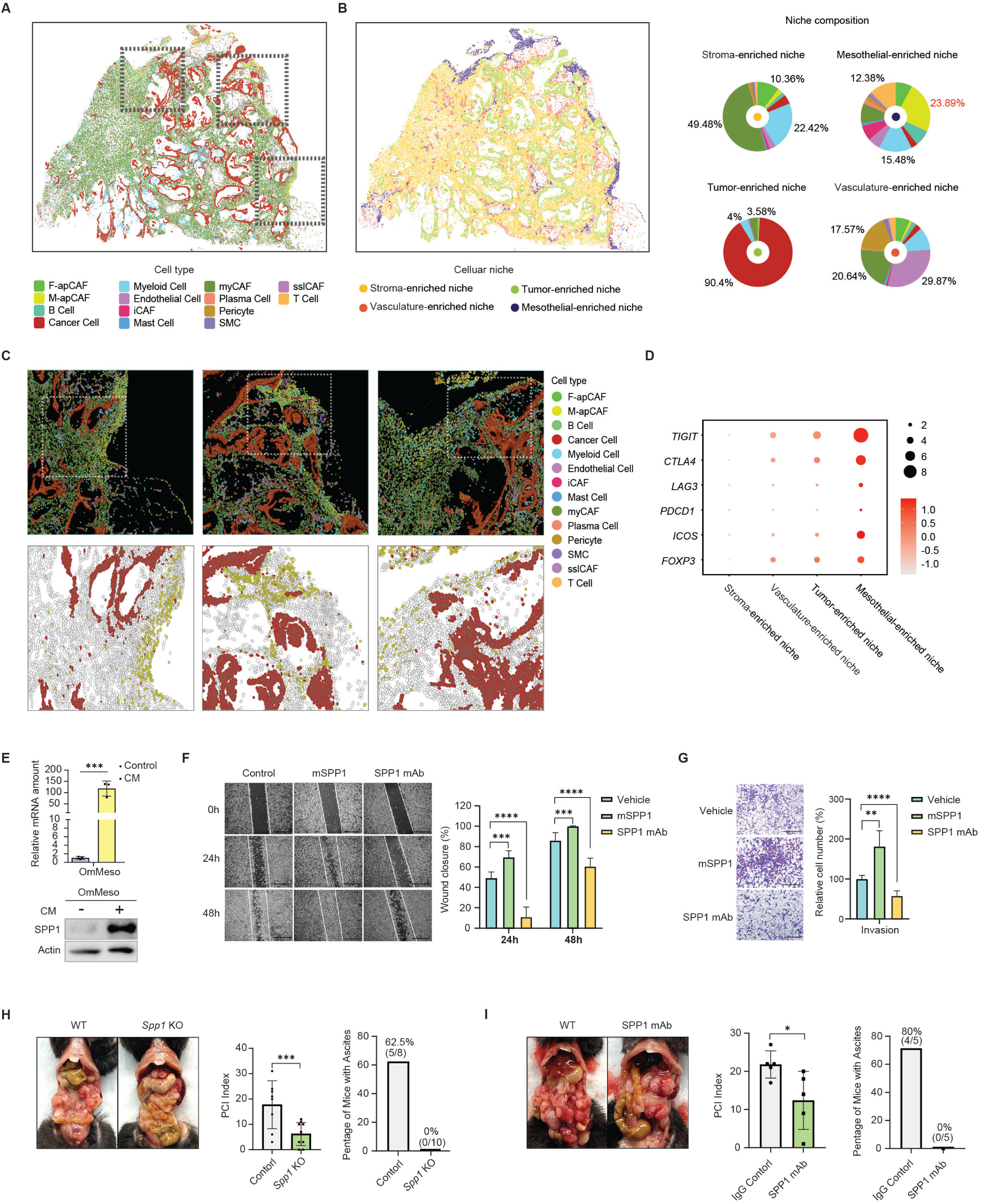
Spatial analysis of M-apCAF niches. (A) Robust cell type decomposition is performed in human PM sample with robust cytokeratin^+^ CAF formation to deconvolve the Xenium data into cell types using our pan-cancer scRNA-seq atlas as reference. (B) Four spatial niches are identified. Percentages of cell types within each niche are shown. (C) Visualization of the spatial distribution of different cell types in four M-apCAF-enriched regions. (D) Expression of T cell immunosuppressive genes across four spatial niches. (E) RT-PCR (n=3/group) and western blots measuring the expression of SPP1 in OmMeso cells after tumor conditioned medium treatment. (F) Wound healing assays are performed to measure the migration capability of MC38 colon cancer cells in the presence of mouse recombinant protein or anti-SPP1 mAb. Representative pictures of cell migration at 0h, 24h, 48h are shown. n=3/group. (G) Matrigel transwell assays in the presence of mouse recombinant protein or anti-SPP1 mAb for 24 hours are performed. Representative pictures for each group are shown. n=3/group. (H) MC38 cancer cells are injected intraperitoneally into wildtype (WT) or *Spp1* knockout (KO) mice on a C57BL/6 background (WT, n=8; KO, n=10). Mice are sacrificed 4 weeks after cancer cell injection. Peritoneal cancer index (PCI) scores and ascites formation are measured. (I) MC38 cancer cells are injected intraperitoneally into wildtype C57BL/6 mice. Mice are treated with control Ab or anti-SPP1 mAb (n=5/group) one week after cancer cell injection and maintained at two doses/week. Mice are sacrificed 4 weeks after cancer cell injection. PCI scores and ascites formation are measured.

Our prior study demonstrated that M-apCAFs were able to induce the formation of immunosuppressive T cells in murine models (Huang et al., 2022). Consistently, we found T cells within the mesothelial-enriched niches expressed a high level of suppressive markers (*TIGIT*, *CTLA4*, *LAG3*, *PDCD1*, *ICOS*, *FOXP3*) (Figure 5D). Further ligand-receptor interaction analysis suggested that *SPP1* was a major ligand that mediated the interaction between M-apCAFs and immune cells or cancer cells (Figure S5D-E).

### Stromal knockout of *Spp1* results in a reduction of PM and ascites formation

SPP1, also known as osteopontin, is a highly phosphorylated secreted protein that normally functions as extracellular structural protein in bone. In the wound and cancer settings, SPP1 is up-regulated as an acute phase protein (Rangaswami et al., 2006). It is associated with inflammation and fibrotic reaction (Lenga et al., 2008; O’Regan and Berman, 2000; Pardo et al., 2005). However, the function of SPP1 is still not completely understood. In cancer, SPP1 is known to exert multifaceted tumor-promoting functions by promoting cellular plasticity and chemoresistance in cancer cells and serving as an immune checkpoint in T cells, but the effect of SPP1 in PM is unknown (Klement et al., 2018; Nallasamy et al., 2021). Interestingly, we observed that iCMS3 PM tends to have a higher level of CAF-derived SPP1 (Figure S5F), potentially contributing to the peritoneal tropism of this PM subtype. SPP1 can be expressed by cancer cells, myeloid cells and CAFs (Bill et al., 2023; Wang et al., 2023). Since our computational data suggest apCAFs is a major source of SPP1 among other CAF subtypes, we validated this result by treating the mesothelial cells isolated from mouse omentum (OmMeso cells) with tumor conditioned media. The conditioned media treatment resulted in a marked elevation of SPP1 expression in OmMeso cells (Figure 5E).

To examine the effect of SPP1 on cancer cell migration and invasion, we performed the scratch assay and Matrigel invasion assay with the murine colon cancer cells MC38 in the presence of recombinant SPP1 protein or anti-SPP1 monoclonal antibody (mAb). We found the addition of SPP1 protein could significantly facilitate while anti-SPP1 mAb inhibited cancer cell migration and invasion (Figure 5F-G). We then investigated the function of stromal SPP1 on promoting PM formation. We injected the syngeneic MC38 cancer cells intraperitoneally into wildtype or *Spp1* knockout mice. We found the stromal *Spp1* knockout resulted in a substantial reduction of PM and ascites formation (Figure 5H). Furthermore, these phenotypes could be recapitulated by treating the wildtype PM-bearing mice with anti-SPP1 mAb (Figure 5I). Taken together, our data suggest that stromal SPP1 is a major driver for PM and apCAFs may form specific niches that produce SPP1 to facilitate this process.

### M-apCAFs are associated with SPP1-positive niches in PDAC

To explore the apCAF niches in primary tumors, we examined the PDAC model, given the previous studies of apCAFs in PDAC (Elyada et al., 2019; Huang et al., 2022). We performed Xenium assays in 3 treatment-naïve and 3 chemoradiotherapy-treated human PDAC samples. Unsupervised clustering analysis identified 6 major cell types including CAFs, endothelial cells, lymphocytes, myeloid cells, and two populations of cancer cells (cancer cell 1 and cancer cell 2) (Figure S6A-B). We found that after the chemoradiotherapy treatment, tissues became more desmoplastic with a larger CAF proportion, consistent with a recent report (Figure S6C) (Cui Zhou et al., 2022). And interestingly, we found there was a reduction of cancer cell 1 in the chemoradiotherapy group while the ratio of cancer cell 2 remained similar, suggesting cancer cell 2 was the chemoradiotherapy-resistant cancer cell population (Figure S6C). Through differential gene analysis, we found cancer 2 was marked by a high level of *SPP1* expression (Figure S6D).

After cell type deconvolution and niche composition analysis, we found that compared to PM, TLS formation in PDAC was relatively rare. Among the six samples, we only found one sample that carried TLS structures. Four niches were identified in this sample including tumor-, stroma-, lymphocyte- and vasculature-enriched niches (Figure 6A and Figure S6E). Moreover, we found the two cancer cell subtypes forming SPP1^-^ and SPP1^+^ tumor epithelium across the tissue. Therefore, we classified the whole tissue into SPP1^-^ cancer, SPP1^+^, stroma and TLS areas and tried to compared the distribution of both apCAF lineages among these areas (Figure 6B-C). We found that the distribution of F-apCAFs was consistent with PM, mostly locating within the TLS regions (Figure 6D). Intriguingly, M-apCAFs were enriched in the SPP1^+^ tumor areas (Figure 6D). Furthermore, we found these areas were infiltrated with SPP1^+^ myeloid cells (Figure 6C), which defines the polarization of a protumor myeloid cell population (Bill et al., 2023). These data suggest that M-apCAFs may contribute to a SPP1^+^ niche that promotes tumor progression and therapy resistance in PDAC.

**Figure 6.**
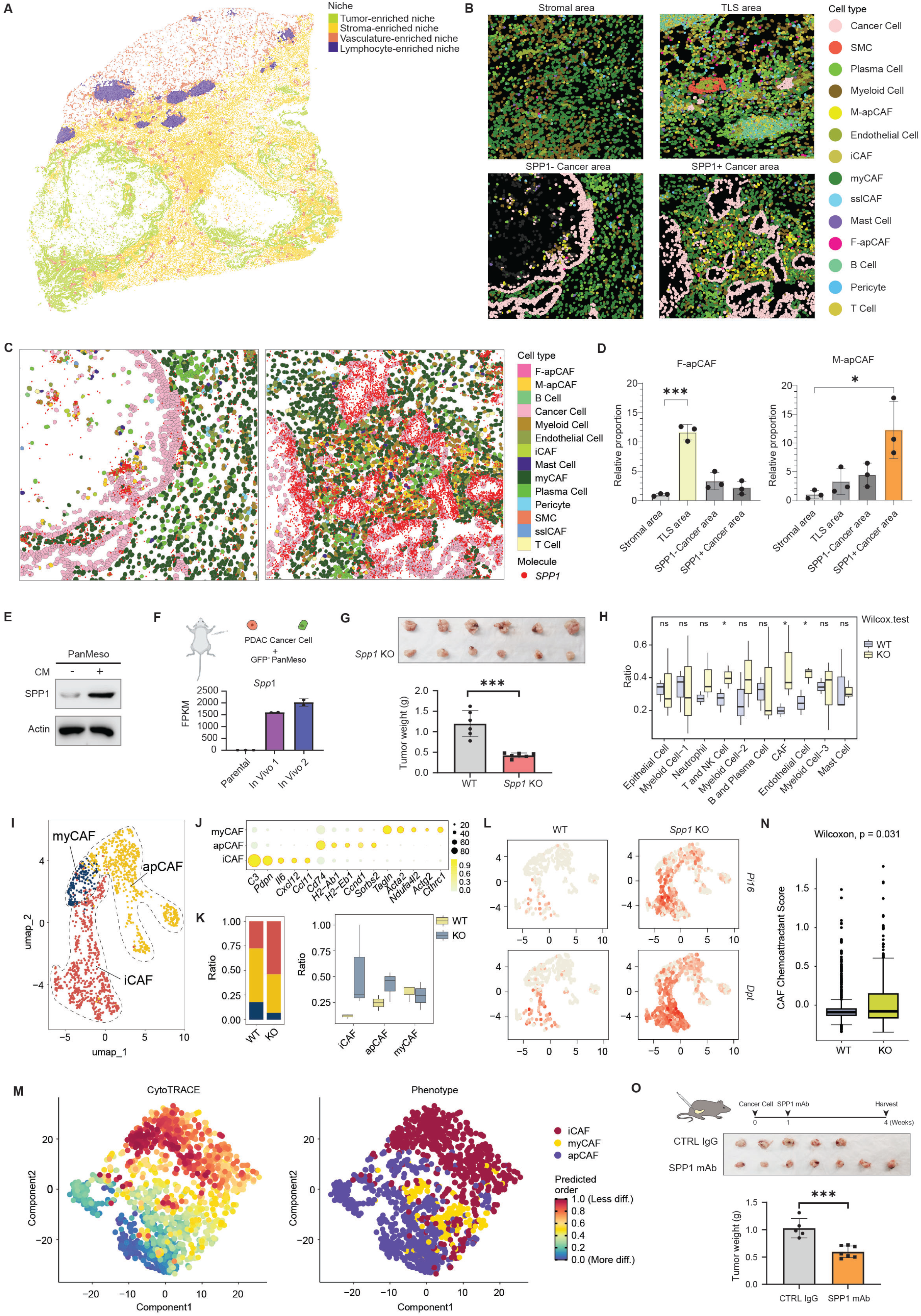
apCAFs contribute to SPP1^+^ tumor-promoting niches. (A) Spatial niches are identified in human PDAC sample with TLS formation. (B) Based on the spatial niches and expression of *SPP1* in cancer cells, PDAC sample is classified into four areas: stroma, TLS, SPP1^-^ and SPP1^+^ cancer. Deconvolved cell types are shown in each area. (C) Expression of *SPP1* is visualized in SPP1^-^ and SPP1^+^ cancer areas. (D) Proportions of F-apCAFs and M-apCAFs are quantified in stromal, TLS, SPP1^-^ and SPP1^+^ cancer areas (n=3 for each area). (E) Western blots measuring the expression of SPP1 in PanMeso cells after tumor conditioned medium treatment. (F) GFP^+^ PanMeso cells are co-injected with a murine PDAC cell line (BMFA3: In Vivo 1 or CT1BA5: In Vivo 2) at a 1:1 ratio. Tumors are harvested 1 month after injection and digested into single-cell suspension. GFP^+^ cells are collected by flow sorting and subjected to RNA-seq analysis in comparison to parental PanMeso cells to evaluate the *Spp1* expression. (G) Syngeneic PDAC cancer cells (6620c1) are injected orthotopically into wildtype (WT) or *Spp1* knockout (KO) C57BL/6 mice (n=6/group). Tumors are harvested 1 month after injection. (H) *Spp1* WT or KO tumors are digested into single cell suspension and subjected to scRNA-seq (6 tumors/group, every two tumors are pooled together for library construction). Ratio of each cell type between WT and KO group is compared and quantified. (I) CAFs from both *Spp1* WT or KO tumors are extracted from the scRNA-seq data. iCAF, myCAF and apCAF clusters are identified. (J) Signature genes of each CAF subtype. (K) Proportional changes of CAF subtypes between *Spp1* WT and KO group. (L) UMAPs showing sslCAF marker *Pi16* and *Dpt* expression between *Spp1* WT and KO tumors. (M) CytoTRACE analysis determining the progenitor and differentiation status among iCAFs, myCAFs and apCAFs, with higher score indicating more stem-like and less differentiated status. (N) Quantification of the expression of T cell chemoattractant genes in CAFs between *Spp1* WT and KO tumors. (O) Syngeneic PDAC cancer cells (6620c1) are injected orthotopically into wildtype C57BL/6 mice. Mice are treated with control Ab (n=5) or anti-SPP1 mAb (n=7) one week after cancer cell injection and maintained at two doses/week. Mice are sacrificed 4 weeks after cancer cell injection.

### Stromal SPP1 facilitates fibrotic barrier formation and T cell exclusion in PDAC

To confirm M-apCAFs were the major source of SPP1 in PDAC, we first treated the pancreatic mesothelial cell line (PanMeso cells) with conditioned media collected from murine PDAC organoids. We found the treatment resulted in an up-regulation of SPP1 expression in PanMeso cells (Figure 6E). To investigate whether the in vivo tumor microenvironment could induce *Spp1* expression in PanMeso cells, we orthotopically co-injected GFP-labeled PanMeso cells with two cancer cell lines derived from genetically engineered mouse model (GEMM) of PDAC respectively (BMFA3 or CT1BA5). Tumors were harvested and digested four weeks after injection. GFP^+^ PanMeso cells were flow sorted and subjected to transcriptomic analysis. We found that GFP^+^ PanMeso cells sorted from the tumors had a drastic increase of *Spp1* expression compared with parental cells (Figure 6F).

We then further investigated the effect of stromal SPP1 on PDAC progression in vivo. A syngeneic PDAC GEMM cell line was injected orthotopically into wildtype or *Spp1* knockout mice. The stromal depletion of *Spp1* led to a significant reduction of tumor growth (Figure 6G). To understand the microenvironmental changes caused by *Spp1* knockout, we digested the tumors from the control and knockout groups into single cell suspension and performed scRNA-seq (Figure S6F-G). The scRNA-seq data suggested a significant increase of T cell infiltration into the stromal *Spp1* knockout tumors (Figure 6H). We confirmed the T cell infiltration by IHC staining (Figure S6H).

However, we also found there was an increase of fibroblasts after *Spp1* knockout (Figure 6H). Therefore, we speculated there might be a phenotypic alteration in fibroblasts that promote T cell infiltration. We then extracted all fibroblasts in the scRNA-seq data from both control and knockout groups and performed unsupervised re-clustering. We were able to identify the presence of iCAFs, myCAFs and apCAFs (Figure 6I-J and Figure S6I). We found the expression of *Spp1* was mostly enriched within the apCAF lineages (Figure S6I). When we compared the proportions of CAF subtypes, we found that there was a reduction of myCAFs, which is the major cause of stromal barrier, and increase of iCAFs in the *Spp1* knockout tumors (Figure 6K) (Krishnamurty et al., 2022). Importantly, CAFs in the knockout group also up-regulated the sslCAF markers such as *Pi16* and *Dpt* (Figure 6L), suggesting after *Spp1* knockout, CAFs became normalized and carried stead-state features found in normal tissues (Figure 1). CytoTRACE analysis further supported iCAFs/sslCAFs were the progenitor state of myCAFs (Figure 6M) (Gulati et al., 2020). We also evaluated the expression of chemoattractants in CAFs that were responsible for T cell infiltration and found the chemoattractant signature was up-regulated in the *Spp1* knockout group (Figure 6N). The normalization of CAFs and up-regulation of chemoattractants may contribute to the increase of T cell infiltration in the *Spp1* knockout tumors. To investigate the therapeutic potential of targeting SPP1 in PDAC, we treated the syngeneic orthotopic tumors with anti-SPP1 mAb, which resulted in a significant reduction of tumor progression (Figure 6O). Altogether, we demonstrated apCAFs were a major source of stromal SPP1, which might facilitate the protumor desmoplasia and inhibit T cell infiltration.

## Discussion

Although various signatures and functions have been reported, CAFs had been traditionally considered as a uniform population of cells in the tumor microenvironment. Due to the breakthrough of single cell transcriptomic and proteomic techniques, laboratories started to use the techniques to profile the CAF heterogeneity in different cancer types, using tissues derived from human or animal models. Many subtypes of CAFs have been reported as a result (Cords et al., 2024; Dominguez et al., 2020; Elyada et al., 2019; Hosein et al., 2019; Hutton et al., 2021; Steele et al., 2020). However, due to the lack of consensus on nomenclatures and subtype markers, it is unclear about the relationship of the CAF subtypes reported by different studies to date (Sahai et al., 2020).

Therefore, large data cross-tissue comparison of CAFs is urgently needed to establish a framework for our understanding of CAFs, similar as a previous study that compares perturbed-state human fibroblasts by Buechler et al. (Buechler et al., 2021). In fact, a recent study by Foster et al. that cross-compared skin, breast and pancreatic cancer found that CAF phenotypes were more conserved that expected and fell into three superclusters: steady state-like, mechanoresponsive, and immunomodulatory CAFs (Foster et al., 2022). This classification system is mostly consistent with the three major subtypes of CAFs, with myCAFs being mechanoresponsive while iCAFs and apCAFs being immunomodulatory. Here, we extended such comparison to 14 types of solid tumors and found iCAFs, myCAFs and apCAFs existed in all tumor types. Moreover, another CAF phenotype that carries the universal fibroblast progenitor signatures reported by Buechler et al. (marked by *DPT*, *PI16*, *CD34*) also contributes to a major proportion of CAFs within the tumor stroma. This CAF population is potentially associated with more benign tumor regions and consistent with the steady state-like supercluster classification. It was also reported by Krishnamurty et al. that under the TGFβ signal, DPT^+^ fibroblast can differentiate into the LRRC15^+^ myCAF population and facilitate immune evasion (Krishnamurty et al., 2022).

By performing scRNA-seq analysis in murine models, Elyada et al. and our group first reported the apCAF population in PDAC (Elyada et al., 2019; Hosein et al., 2019). apCAFs are defined by their expression of MHC II genes (human leukocyte antigens or HLA genes in human), such as *CD74* and *HLA-DR*. apCAFs have been detected in different cancer types (Costa et al., 2018; Cui Zhou et al., 2022; Sun et al., 2023; Varveri et al., 2024). However, due to the fact that they are less abundant than myCAFs and more context dependent and their cells of origin are incompletely understood, there is still debate whether apCAF is a true CAF population (Hwang et al., 2022; Werba et al., 2023). Moreover, multiplex staining of apCAFs remains difficult because most of the staining relies on MHC II molecules, which are widespread in the tumor microenvironment. Therefore, there is an urgent need for understanding the lineages of apCAFs by big data cross-tissue analysis and applying spatial transcriptomic at a single cell resolution to identify their spatial niches. Here, we found apCAFs were present in the majority of solid tumors. Surprisingly, we found apCAFs were derived from two distinct lineages. One lineage which we call M-apCAF is associated with mesothelial-like cells as we reported previously (Huang et al., 2022). Mesothelial cells carry both epithelial and mesenchymal features and are plastic under certain circumstances (Koopmans and Rinkevich, 2018). In addition, we found this lineage expressed many cytokeratin genes. Therefore, we speculate other mesothelial-like epithelial cells can also serve as progenitors for M-apCAFs in a tissue-dependent manner. Another lineage of apCAFs is associated with fibrocytes and thus we call it F-apCAF. Fibrocytes are bone marrow-derived cells that have signatures of both fibroblasts and myeloid cells and can produce extracellular matrix and inflammatory proteins (Direkze et al., 2004; Phillips et al., 2004). Despite their low abundance in circulation, they strongly promote inflammation and fibrosis by migrating to injury sites.

Given apCAFs carry signatures also expressed by immune cells, dissecting the accurate spatial information of apCAFs requires a spatial transcriptomic platform with single cell resolution. Here we exploited the Xenium high-plex single cell in situ spatial imager. To ensure the rigor of the spatial analysis, we fully customized the whole 480-gene panel of probes based on the cell type markers identified from our integrated cell atlas. This approach enabled us to deconvolve the spatial information using the cell atlas as reference. We then profiled the PM and PDAC patient tissues with Xenium platform, revealing two niches formed by the two lineages of apCAFs: F-apCAFs localize in the TLS areas while M-apCAFs tend to be adjacent to tumor epithelium. Fibroblast is a key orchestrator for lymphoid organ structure and function such as spleen and lymph nodes (Golub et al., 2018). However, fibroblasts involved in TLS formation are poorly understood. Given the proximity of F-apCAFs with TLS, they may play a critical role for TLS formation. This is in line with the fact that fibrocytes have been reported to be associated with an inflammatory phenotype (Reilkoff et al., 2011). In addition, in a study reported by Kerdidani et al., they found apCAF promoted tumor immunity by C1q in lung cancer (Kerdidani et al., 2022). Given C1q is produced by F-apCAFs, it is likely that the results from this study are due to the presence of F-apCAFs. In contrast, M-apCAFs are commonly found adjacent to cancer cells. The formation of M-apCAF niches is more context-dependent. Since mesothelial-like cells are less likely to be recruited into tumor stroma from circulation as fibrocytes, the formation of M-apCAF probably relies on the presence of the progenitors and local differentiation signal. For example, in PM, we found these niches are more robust in areas near the mesothelium regions. Myeloid cells and T cells are also present in the M-apCAF niches. however, we found these T cells carried exhausted and immunosuppressive phenotype. Therefore, M-apCAFs may drive immunosuppression, consistent with our previous study (Huang et al., 2022).

One of the most up-regulated genes in both apCAF lineages is *SPP1*. SPP1 is an acute phase protein that is expressed in a wound setting. It has been reported to be associated with fibrosis, inflammation, immunosuppression and cell stemness (Klement et al., 2018; Nallasamy et al., 2021). Therefore, it might be a critical protein that has multiple functions for promoting wound healing and regeneration. SPP1 is known to be expressed by cancer cells and stromal cells. A recent study has reclassified macrophages based on SPP1 expression to represent the protumor macrophages instead of the classic M1-M2 system (Bill et al., 2023). Here, we validated apCAFs as a major source of stromal SPP1. We also found that stromal knockout of *Spp1* could significantly reduce the PM and primary PDAC progression. Moreover, our data showed targeting SPP1 may be a promising therapeutic strategy for PM and PDAC. This is important and clinically feasible, given many humanized SPP1 antibodies are already in clinical trials for other diseases (Farrokhi et al., 2018).

We have identified two subtypes of PM (iCMS2 and iCMS3) in colon cancer, characterized by differences in epithelial signatures and stromal niches. iCMS3 PM favors peritoneal spread and induces robust TLS and apCAF formation. The strong association between the iCMS3 phenotype and PM allows better stratification strategy in prospective clinical trials. Furthermore, characterizing this subtype may uncover the mechanisms of peritoneal progression and offer novel therapeutic targets leveraging its tumor microenvironment. The lack of effector activity of infiltrating T cells strongly suggests immunosuppressive signals exists. One of such signals may come from apCAFs. Therefore, future studies should evaluate the combinatory effects of tools that target apCAF signal such as anti-SPP1 Ab and immune checkpoint blockade. Nallasamy et al. reports that SPP1 drives cancer cell stemness in PDAC (Nallasamy et al., 2021). We found that SPP1^+^ cancer cells are resistant to chemoradiotherapy and M-apCAFs contribute to the SPP1-enriched niche formation. Therefore, effects of targeting SPP1 to overcome chemoradiotherapy resistance in PDAC should be explored clinically.

In summary, our study provides a robust cross-tissue comparison of apCAFs and identifies two distinct apCAF lineages, one derived from fibrocytes, another from mesothelial-like cells. We characterize the spatial niches of the two apCAF lineages at an unprecedented resolution. These data greatly advance our current understanding of this CAF subtype.

## Acknowledgements

This work was supported by NIH R00 CA252009 to H.H. We acknowledge input from members of the Brekken laboratory. We thank UT Southwestern Core facilities supported in part by the Cancer Center Support Grant (P30 CA142543). Core facilities used included the Tissue Management Core, the Microarray and Immune Phenotyping Core, the Histo Pathology Core, the Flow Cytometry Core and the Whole Brain Microscopy Facility.

## Author contributions

XC: acquisition of data, analysis, interpretation of data and drafting of the manuscript. ZZ: acquisition of data. ZY: acquisition of data. LX: acquisition of data. FR: acquisition of data. YL: acquisition of data. BZ: acquisition of data. PMP: tissue collection. HJZ: interpretation of data. ACK: study concept and design, tissue collection and interpretation of data. HH: study concept and design, acquisition of data, analysis, interpretation of data and drafting of the manuscript.

## Declaration of interests

The authors declare no competing interests.

## Methods

### Resource availability

#### Lead contact

Further information and requests for resources should be directed to and will be fulfilled by the lead contact, Huocong Huang

## Materials availability

All unique reagents generated in this study are available from the lead contact with a completed Materials Transfer Agreement.

## Data and code availability

Details of the published scRNA-seq datasets we used for integrated analysis can be found in the supplementary table. The spatial transcriptomic data and scRNA-seq generated in this study have been uploaded to GEO under the following accession numbers GSE274609, GSE274623, GSE274673. RNA-seq data of PanMeso cells have been previously deposited at GEO (GSE196740). Additional information related to the data in this study is available from the lead contact upon request.

## Experimental model and subject details

### Cell lines

BMFA3 and CT1BA5 were mouse primary pancreatic cancer cell lines derived from *Kras^LSL-G12D/+^*;*Trp53^fl/fl^*;*Pdx1^Cre/+^*(*KPfC*) mice on a C57BL/6 background and isolated as described previously (Huang et al., 2019). *Kras^LSL-G12D/+^*;*Trp53^LSL-R172H/+^*;*Ptf1a^Cre/+^*(*KPC*) cell clone 6620c1 on a C57BL/6 background was purchased (Kerafast) and has been published in a previous study (Li et al., 2018). MC38 murine colon adenocarcinoma cells on a C57BL/6 background were purchased (MilliporeSigma). PanMeso and OmMeso cells were established as follows. Mesothelium was harvested from normal pancreata or omentum of immortomice under the dissecting microscope. The pancreatic or omental mesothelium was seeded onto a tissue culture dish and cultured at 33°C for 20 days. Once cells became confluent, omental mesothelial cells were collected and named omental mesothelial (OmMeso) cells while pancreatic mesothelial cells were subjected to cell sorting. Podoplanin^+^MHC II^+^ cells were collected and named pancreatic mesothelial (PanMeso) cells. All mouse cancer cell lines were cultured in DMEM (Invitrogen) containing 10% FBS and 1% penicillin/streptomycin (Gibco/Thermo) and maintained at 37 °C in 5% CO_2_. The PanMeso and OmMeso cells were cultured in the mesothelial cell media (medium 199 (Gibco/Thermo), 10% FBS, 1% penicillin/streptomycin (Gibco/Thermo), 400 nM hydrocortisone (MilliporeSigma), 870 nM zinc-free bovine insulin (MilliporeSigma), 20 mM HEPES (Thermo Fisher Scientific)), maintained at 33 °C, washed with PBS three times and cultured in new mesothelial cell media at 37 °C before any experiments. All cell lines in this study were confirmed to be free of mycoplasma (e-Myco kit, Boca Scientific) before use.

### Animals

6-week-old male and female mice were purchased from The Jackson Laboratory, including C57BL/6J mice (JAX stock #000664), *Spp1* knockout mice (JAX stock #004936) and immortomice (JAX stock #032619). All animals were housed in a pathogen-free facility with 24-hr access to food and water. Animal experiments in this study were approved by and performed in accordance with the institutional animal care and use committee at the UTSW Medical Center at Dallas. Mice were euthanized by cervical dislocation under anesthesia.

### Patient samples

Formalin-fixed paraffin-embedded (FFPE) tissue sections of T4 colon cancer, PM (from patients who had undergone cytoreductive surgery for colon cancer-derived PM), PDAC (treatment naïve and chemoradiotherapy treated) were obtained with voluntary patient consent from UTSW Tissue Management Shared Resource, which were obtained under a protocol approved by the UTSW Institutional Review Board. All specimens were reviewed by a board-certified pathologist specializing in gastrointestinal pathology to confirm diagnosis. FFPE blocks were cut into 5-μm-thick serial sections. One section was stained with hematoxylin and eosin for pathology review. Tissue sections were used for IHC or spatial transcriptomics in this study. Age and gender of the patients were undisclosed.

## Method details

### Plasmids

pLV-eGFP (Addgene #36083) was used to expressed GFP in PanMeso cells.

### Immunohistochemistry

FFPE tissues were cut in 5-μm sections. Sections were evaluated by single color or multiplex immunohistochemical analysis following our previously reported protocol using antibodies specific for SPINK4 (Invitrogen, 1:500), CD3 (Invitrogen, 1:300), CD8 (for human, Abcam, 1:300; for mouse, Cell Signaling, 1:300), FoxP3 (R&D Systems, 1:500), pan-cytokeratin (Thermo Fisher Scientific, 1:500) (Sorrelle et al., 2019). For brightfield images, slides were scanned by NanoZoomer 2.0-HT digital slide scanner (Hamamatsu) and images were obtained with NDP.view2 software (Hamamatsu) and analyzed by ImageJ. For fluorescent images, images were obtained with ECHO Revolve microscope and analyzed by ImageJ.

### RT-PCR

RNA was extracted using RNeasy Mini Kit (Qiagen). cDNAs were synthesized using the iScript cDNA synthesis kit (Bio-Rad). The expression of genes was measured by qRT-PCR with SYBR-Green Master Mix (Bio-Rad).

### Western blot analysis

Cells were lysed using a radioimmunoprecipitation assay buffer (50 mM Tris-HCl, pH 8.0, 150 mM NaCl, 0.1% SDS, 0.5% sodium deoxycholate, and 1% Nonidet P-40) for SDS-PAGE. Protein concentrations were measured with a BCA Protein Assay Kit (Thermo Fisher Scientific). Laemmli sample buffer was added to the lysates, which were then boiled for 10 minutes. Proteins were separated by SDS-PAGE, transferred to nitrocellulose membranes, and blocked with 5% nonfat dry milk or bovine serum albumin. After blocking, membranes were incubated for 1 hour with primary antibodies against SPP1 (Bio X Cell, 1:1000) in TBST (10 mM Tris-HCl, pH 7.5, 150 mM NaCl, 0.05% Tween 20). The membranes were then washed three times for 10 minutes each with TBST, followed by a 1-hour incubation with horseradish peroxidase-conjugated anti-mouse or rabbit secondary antibodies (Jackson ImmunoResearch Laboratories). After secondary antibody removal with three 10-minute TBST washes, the membranes were treated with SuperSignal West Pico substrate (Thermo Fisher Scientific) to visualize the immunoreactive bands.

### Tumor conditioned medium

Mouse PDAC organoids were obtained from *KPfC* tumors. The tumor tissues were finely chopped and incubated at 37°C for 12 hours in a digestion buffer, which included 0.012% collagenase XI (MilliporeSigma) and 0.012% dispase (Gibco) in DMEM (Invitrogen) with 1% FBS. The resulting organoids were then pelleted, rinsed with PBS, and plated on a Petri dish in DMEM (Invitrogen) with 10% FBS. After a 4-day culture period of PDAC organoids or MC38 cancer cells, the medium was passed through a 0.22 μm ultra-low protein binding filter and collected as PDAC or colon cancer conditioned medium.

### Wound healing assay

For the wound healing assay, cells were plated onto 6-well tissue culture plates coated with 50 μg/ml Matrigel (BD Biosciences) with or without 100 ng/ml recombinant mouse SPP1 protein (R&D Systems) or 1 μg/ml SPP1 monoclonal antibody (Bio X Cell). After the cell monolayer reached confluency, the monolayer was scraped with a 200 μl tip to generate scratch wounds. The cells were washed with PBS and then cultured in medium containing 10% FBS in a tissue culture incubator. At the designated time points after scratch wounds, plates were washed with PBS, and the wound width was photographed and measured under ECHO Revolve microscope.

### Matrigel invasion assay

For the Matrigel invasion assay, transwells were precoated with 100 ng/ml recombinant mouse SPP1 protein (R&D Systems) or 1 μg/ml SPP1 monoclonal antibody (Bio X Cell), with or without 150 μg/cm² Matrigel (BD Biosciences). 5×10⁴ MC38 cells were added to the transwells and allowed to migrate towards medium with 10% FBS. After 24 hours, the cells on the top side of the transwells were scraped off with cotton swabs. The transwell membrane was then fixed with 4% paraformaldehyde for 20 minutes. Cells were stained with crystal violet and counted under microscopy at 100× magnification.

### In vivo models

For PM, MC38 cells were injected intraperitoneally (5 × 10^4^ cells) in 6-week-old C57BL/6 mice (male and female). Mice were sacrificed 1 month after injection and PCI score was calculated based on published literature (Lenos et al., 2022). For PDAC, 6620c1 cells were injected orthotopically (2 × 10^5^ cells) in 6-week-old C57BL/6 mice (male and female). To harvest tumors, mice were sacrificed 1 month after injection. Tumors were then subjected to digestion, scRNA-seq or IHC staining. For therapies, C57BL/6 mice were treated with control (clone MOPC-21, BioXCell, 500 μg/dose) or anti-SPP1 Ab (MPIIIB10, BioXCell, 500 μg/dose) 7 days after cancer cell injection and maintained two doses per week during the tumor formation.

### Single cell cDNA library preparation and sequencing

Single cells were isolated from mouse tumor tissues following a previously established method. Initially, a 10x digestion buffer was prepared in PBS, containing collagenase type I (450 units/ml, Worthington Biochemical), collagenase type II (150 units/ml, Worthington Biochemical), collagenase type III (450 units/ml, Worthington Biochemical), collagenase type IV (450 units/ml, Thermo Fisher Scientific), elastase (0.8 units/ml, Worthington Biochemical), hyaluronidase (300 units/ml, MilliporeSigma), and DNase type I (250 units/ml, MilliporeSigma). Freshly dissected tumor tissues were cut into 1-2 mm³ pieces using a sterile blade, washed with PBS, and then incubated in 50 mL of 1x digestion buffer on a shaker at 37°C for 45 minutes. The resulting cell suspension was filtered through a 40 μm mesh (MilliporeSigma) and centrifuged at 400 g for 5 minutes. After removing the supernatant, the cell pellet was resuspended in ACK lysing buffer (Thermo Fisher Scientific) and incubated on ice for 5 minutes to lyse red blood cells.

For each group, we included 6 tumors. Every two tumors were then pooled together for library construction (three libraries per group). Approximately 10,000 cells were used for library construction with the Chromium Next GEM Single Cell 3′ Kit v3.1 (10x Genomics), following the manufacturer’s protocol. The purified libraries were sequenced on an Illumina NextSeq6000. Raw base call (BCL) files from the sequencer were demultiplexed into fastq files. These fastq files were aligned to the mm10 mouse reference genome and quantified using the Cell Ranger pipeline (version 7.2.0, 10x Genomics). The preliminary filtered data generated from Cell Ranger were utilized for all subsequent analyses.

### Single-cell RNA-seq data processing

The count matrices generated by Cell Ranger were imported using the Read10X function from the Seurat package (v5.0.1) and converted into Seurat objects. Quality control was performed based on the number of detected genes and the proportion of mitochondrial gene counts per cell. Specifically, cells with fewer than 200 detected genes or more than 6000 detected genes, as well as cells with a mitochondrial unique molecular identifier (UMI) count percentage greater than 15%, were excluded. Additionally, genes detected in fewer than 3 cells were removed. DoubletFinder (v2.0.3) with default settings was used to identify and exclude potential doublets.

Normalization was carried out using the NormalizeData function to ensure uniform total gene expression per cell, with the scaling factor set to 10,000. The FindVariableFeatures function identified the top 2000 highly variable genes (HVGs). Seurat objects were then scaled with the ScaleData function before performing principal component analysis (RunPCA), retaining the first 50 principal components (PCs) for further analysis. Seurat objects from multiple samples were merged using the merge function. To correct for batch effects, the Harmony algorithm (v1.2.0) was applied with default parameters (RunHarmony, max.iter.harmony = 10) using sample labels as covariates.

A shared nearest neighbor (SNN) graph was constructed using the FindNeighbors and FindClusters functions in Seurat, leveraging the appropriate PCA dimensions. Cells were clustered using the Louvain algorithm in a graph-based approach. The RunUMAP function (reduction = "harmony") was used to generate the UMAP embedding. Marker genes and differentially expressed genes for each cluster were identified using the FindAllMarkers and FindMarkers functions in Seurat. Cell classification into major lineages was guided by literature marker genes. Visualization was achieved using the DimPlot, FeaturePlot, and VlnPlot functions.

Subclustering analysis followed a similar methodology, including normalization, identification of variable features, dimensionality reduction with Harmony-based batch correction, and cluster identification.

### Cell cluster proportion and hierarchical clustering of different cancer types

To quantify the distribution of cell clusters across various cancer types, we calculated the proportion of each cell cluster within each cancer type. Additionally, we performed unsupervised hierarchical clustering for all cancer types based on these cluster proportions. The frequencies of all clusters served as input for the hierarchical clustering, which was carried out using the hclust function.

### Calculation of signature score

To assess the functional differences among CAF subpopulations, we identified gene sets corresponding to 10 specific gene signatures (Table S3). The AUCell package (v1.24.0) in R was used to compute the signature scores for these gene sets. Initially, the AUCell_buildRankings function was utilized to create the ranking expression matrix. Next, the AUC values were calculated using the AUCell_calcAUC function. Visualization of the results was accomplished with the ggplot2 package in R.

### Similarity analysis of CAF subclusters

To measure the similarity between different CAF subclusters, we utilized the AverageExpression function from the Seurat package to compute the average expression values for all genes across the subclusters. The correlation coefficients between the subclusters were then calculated using the corr.test function from the psych package (v2.4.3). Visualization was performed with the pheatmap package (v1.0.12).

### Enrichment of fibroblast subclusters in normal and tumor tissues

To quantify the enrichment of each fibroblast subcluster in normal and tumor tissues, we initially grouped all cells by tissue types and cell subclusters to determine the total number of cells in each group, converting this information into a matrix. We then computed the marginal tables for rows, columns, and the overall dataset, followed by normalizing the data based on these marginal tables. Tissue types with normalized values greater than 1 were categorized as enrichment, while those with values less than 1 were categorized as depletion.

### Pseudotime trajectory inference

Monocle3 (v2.30.1) was used to infer cell type transition states among apCAF subclusters. Dimensionally-reduced data from the integrated Seurat object was converted into Monocle3 cell_data_set objects. To facilitate the learning of disjoint graphs across multiple partitions, cells were re-clustered. A trajectory graph was then constructed, accommodating closed loops and multiple partitions. For each compartment, the starting point of the trajectory was determined as the node closest to the previously identified progenitor populations. Pseudotime along the trajectory was calculated using this designated starting node.

### Gene regulatory network inference

The SCENIC algorithm in Python (pySCENIC v0.12.1) was used to infer regulatory interactions between transcription factors and their target genes. The process began by constructing a gene co-expression network using the grn function with the GRNBoost2 algorithm. Regulons were subsequently refined based on the hg38 cistarget database. The aucell command was then employed to score regulon activity in each cell. Differentially activated regulons within each CAF subset were identified using the wilcox.test with BH-adjusted p-values < 0.05. Additionally, ChIP-seq binding peaks of OCT4 at the *SPP1* promoter were visualized using the WashU Epigenome Browser from the Cistrome project.

### Cell-cell interaction analysis

The Python package CellPhoneDB (v3.0) was utilized to infer interactions between apCAFs and immune cells/cancer cells. Input files for the statistical analysis, including the raw count matrix and the cell type annotation files, were extracted from the Seurat object we created. Enriched ligand-receptor pairs between cell subclusters were identified by performing a permutation test on the expression matrix. Significant ligand-receptor pairs with a p-value < 0.05 were selected for illustration.

### Survival analysis

We separated apCAFs into normal and tumor tissues for differential expression analysis, identifying six genes (*NDUFA4L2*, *EGLN3*, *SPP1*, *ANGPTL4*, *PLOD2*, and *HILPDA*) as cancer-related apCAF signature. Survival analysis was subsequently conducted using the GEPIA2 database, based on the expression levels of individual gene or the multi-gene signature, and Kaplan-Meier curves were generated to visualize the results.

### Enrichment analysis

Pathway enrichment analysis was conducted with Metascape (https://metascape.org) or GSEA, and cell-type specific enrichment analysis was performed using WebCSEA (https://bioinfo.uth.edu/webcsea/).

### CytoTRACE analysis

The CytoTRACE package was used to evaluate the transcriptional diversity of each cell, focusing on their stem cell or differentiation status. Each cell was assigned a CytoTRACE score, where higher scores corresponded to greater stemness and less differentiation characteristics.

### GeoMx spatial profiling

Nanostring GeoMx Human Whole Transcriptome Atlas (WTA) platform was used to profile a total of 96 ROIs from 8 PM patient samples (12 per sample). This panel profiles the whole transcriptome by targeting over 18,000 unique transcripts from human protein encoding genes plus ERCC negative controls. The panel excludes uninformative high-abundance RNAs such as ribosomal subunits and includes RNA probes designed for Illumina NGS readout with the Seq Code library prep. Tissue slides were incubated with the fluorophore-conjugated antibodies against vimentin (Santa Cruz, clone E-5), pan-cytokeratin, and a cocktail of photocleavable oligonucleotide probes from the GeoMx kit. A board-certified pathologist selected ROIs based on histology and morphology marker staining. Libraries were prepared according to the GeoMx manual and sequencing depth was 100 read pairs/μm2.

Process sequenced FASTQ files were first converted to DCC (digital count files) run by Nanostring’s GeoMx NGS Pipeline (Version 2.3.3.10). DCCs were transferred to the GeoMx DSP Data Analysis Suite. Readout-specific quality control was checked and standardized QC threshold settings was set as recommended by the manufacturer. All 96 ROIs passed the segment quality control. Gene filtering retained 11903 of 18677 (63.73%) genes after removing targets with expression at or lower than selected threshold value at or above specific frequency (5% in our case). Q3 normalization was applied to account for differences in sequencing yield, enabling biological comparison of gene expression across ROIs. The normalized count density distributions across individual ROIs were grossly similar, and there was no observable bias among samples or by histology (Table S5).

### GeoMx data analysis

To extract the iCMS2 and iCMS3 epithelial signatures, we downloaded the count expression matrix and metadata for epithelial cells from Synapse Storage (ID: syn26844071). For gene expression counts, individual samples were merged and analyzed using the Seurat 4.3 software package. Cells with UMI counts less than 500 or more than 20% mitochondrial counts were removed. Further filtering of cells in the Seurat object was done by removing normal epithelial cells. Epithelial cancer cell spatial deconvolution for each segmented tumor ROI were performed with the Bioconductor package SpatialDecon based on the epithelial cell profile matrix from the iCMS dataset. Principal component analysis for spatial data of epithelial cells were generated by ClustVis. Samples were grouped into iCMS2 or iCMS3 based on the deconvolution information. Top 500 differential expressing genes of each group were used to complete the analysis (Table S6-S7). Differentially expressed genes (DEGs) of GeoMx RNA data between two iCMS cancer cell or iCMS CAF subgroups were determined using the limma package in R. |Log2FC|> 1, and FDR < 0.05 were considered statistically significant. The volcano plots of DEGs were visualized by ggpubr R package.

### Imaging mass cytometry

5-μm-thick sections were cut from FFPE tissue blocks onto glass slides (3 iCMS2 and 3 iCMS3 PM samples) and stained with a cocktail of metal-conjugated antibodies against immune markers as well as cytoplasmic and nuclear markers provided by the company that were used for cell segmentation (Fluidigm). The tissue sections were then ablated by Hyperion system for data acquisition (Fluidigm). Two images per sample were screened and the MCD files were processed with MCD Viewer. TIFF files were then extracted from MCD files for downstream analyses. For cell segmentation, machine learning was utilized in CellProfiler 4.2.5 for pixel classification of cell nuclei, enabling cell segmentation. CellProfiler generates segmentation masks that, when combined with individual TIFF files, allow for the extraction of single-cell information from each image. Next, histoCAT (high-dimensional cytometry analysis toolbox) was employed to extract single-cell data from the images of the segmentation masks. histoCAT enables the measurement of abundance for all markers of interest within cells and regions of interest. This step helped capture the high-dimensional characteristics of each individual cell. Then, we output standardized data from histoCAT and imported the Seurat4.3 R package for further analysis.

### Xenium in situ high-plex assay

Based on the single-cell atlas data, we designed a custom gene panel (Table S4) comprising 480 genes for cell type recognition and gene functional analysis. Tissue sections were prepared according to the Xenium In Situ for FFPE-Tissue Preparation Guide protocol (10X Genomics). Briefly, FFPE sections were cut to 5 μm and placed in the designated sample area (10.45mm x 22.45mm) on a Xenium slide (10X Genomics). The sections were then dewaxed in xylene, rehydrated through a gradient ethanol series, and treated with enzyme to facilitate mRNA accessibility. Probe hybridization was performed at 50 °C overnight, followed by the removal of unhybridized probes. Locked DNA probes bound to target mRNA were ligated at 37 °C for 2 hours (Xenium ligase A/B). Circular DNA probes for each gene underwent rolling circle amplification, producing multiple copies of specific DNA barcodes. After washing, background fluorescence was chemically quenched. The slides were then analyzed using a Xenium analyzer (10X Genomics) for image acquisition. The morphological image of DAPI nuclear staining was used for cell assignment, and built-in machine learning algorithms were applied for cell segmentation. Output files generated by the instrument’s pipeline were used for downstream analysis.

### Xenium data processing

We followed the Seurat vignette (https://satijalab.org/seurat/articles/seurat5_spatial_vignette_2.html) to load and analyze the Xenium output data using the Seurat package. We read and merged the cell x gene matrices to create a Seurat object. The SCTransform function was used for normalization, followed by standard dimensionality reduction and clustering. The clustering results were visualized in UMAP space using the DimPlot function.

For each Xenium sample, we performed robust cell type decomposition (RCTD) with the R package spacexr (v2.2.1) to deconvolve spatial transcriptomic data into cell types, using the Seurat reference constructed from our pan-cancer scRNA-seq atlas data. The cell type deconvolution results were incorporated into the Seurat object metadata for subsequent analysis. The BuildNicheAssay function was used to compute spatial niches by calculating the cell type composition of each cell using the k-nearest neighbor algorithm. Cells with similar neighborhoods were grouped into spatial niches using k-means clustering. Seurat’s ImageDimPlot and ImageFeaturePlot functions were leveraged to visualize transcript localization and segmentation boundaries. Additionally, these visualizations, along with clustering outputs, were produced using the Xenium Explorer 2 software.

Spatial distance analysis between F-apCAFs, myCAFs, and TLS was conducted in the TLS regions. T, B, and Plasma cells were classified as TLS cells. For the F-apCAF or myCAF subset, the distance between a specific cell *k* in this subset and TLS was defined as the shortest Euclidean distance *d* between cell *k* and all cells *j* in the TLS, expressed as:

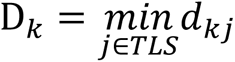

Finally, we calculated the kernel density estimates of all D_*k*_ in a specific subset using the density function and visualized it.

### Quantification and statistical analysis

Statistical analyses and graphical representations of the data were primarily performed using the R statistical environment (https://cran.r-project.org, version 4.3.2) and GraphPad Prism software (version 10.0.0). Relative risks (RR) were computed using bivariate log-binomial regressions, considering different sites of metastasis as the outcome and SPINK4 (+ or -) as the exposure. This analysis was conducted with SAS 9.4 statistical software, with significance determined by a two-sided α level of ≤0.05. The Wilcoxon Rank-Sum test was used to compare means between two groups of continuous variables, while Spearman’s rank correlation was employed to assess correlations across multiple groups. The Benjamini-Hochberg method was applied for multiple test corrections, and false discovery rates (FDR) were calculated accordingly. In vivo studies were quantified with t test. A P or FDR value of less than 0.05 was deemed statistically significant, with symbols for significance as follows: ns, non-significant; *<0.05, **<0.01, ***<0.001, ****<0.0001. Illustrations were created using BioRender.com or Adobe Illustrator software.

---

**Figure S1.**
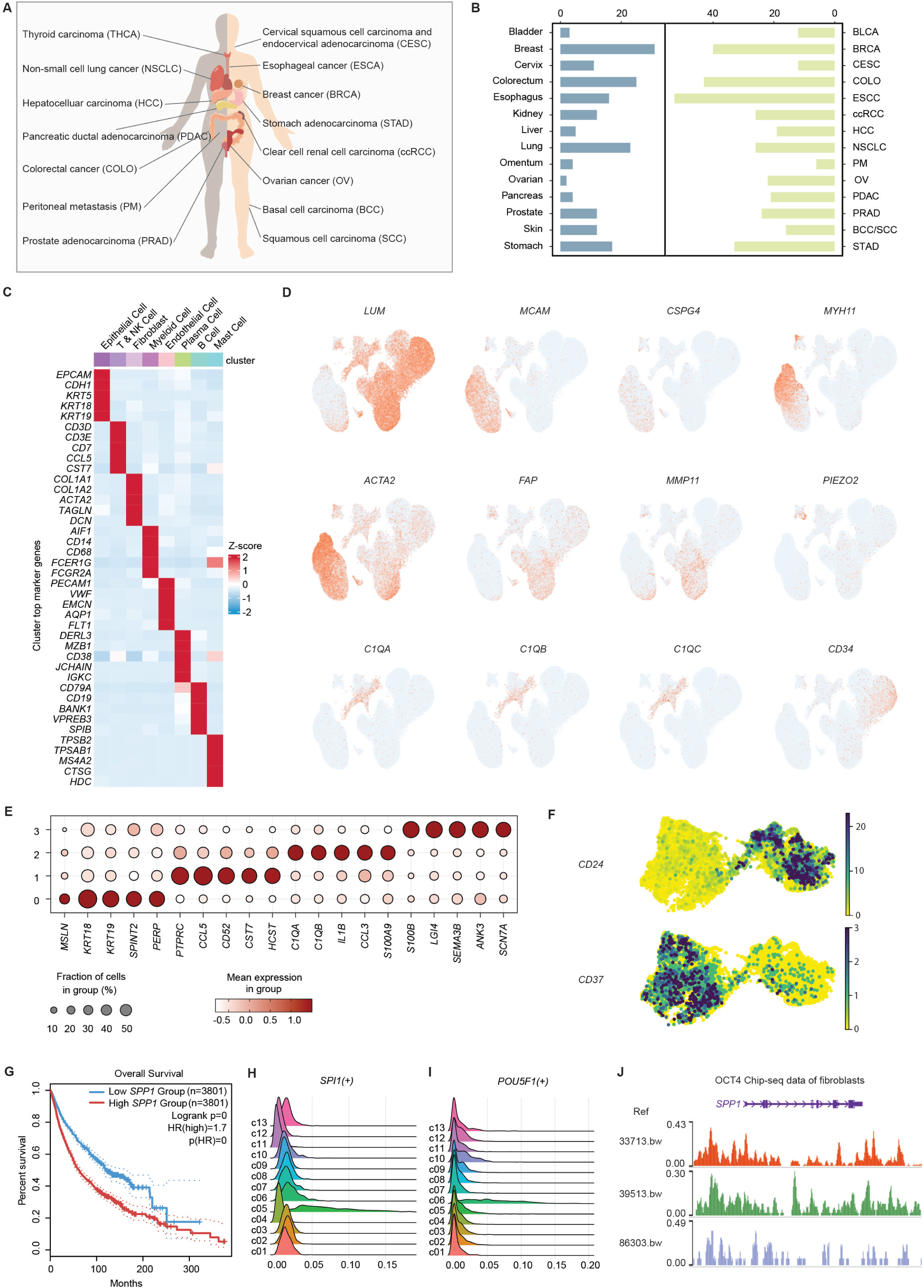
Additional molecular signatures of CAFs in pan-cancer integrated analysis, Related to Figures 1 and 2. (A) Scheme of human cancer types included in the integrated analysis. (B) Numbers of samples in each normal tissue or cancer type. (C) Marker genes of the major cell types in the pan-cancer scRNA-seq atlas. (D) UMAPs showing *LUM*, *MCAM*, *CSPG4*, *MYH11*, *ACTA2*, *FAP*, *MMP11*, *PIEZO2*, *C1QA*, *C1QB*, *C1QC*, *CD34* expression in CAFs. (E) Signature genes of apCAF subclusters. (F) Expression of CD24 and CD37 in the two apCAF lineages. (G) Combined overall survival of the 14 types of cancer with *SPP1* expression. (H) Regulon of *SPI1* in each CAF subcluster revealed by SCENIC algorithm. (I) Regulon of *POU5F1* in each CAF subcluster revealed by SCENIC algorithm. (J) ChIP-seq binding peaks of OCT4 at the *SPP1* promoter in fibroblasts visualized using the WashU Epigenome Browser from the Cistrome project.

**Figure S2.**
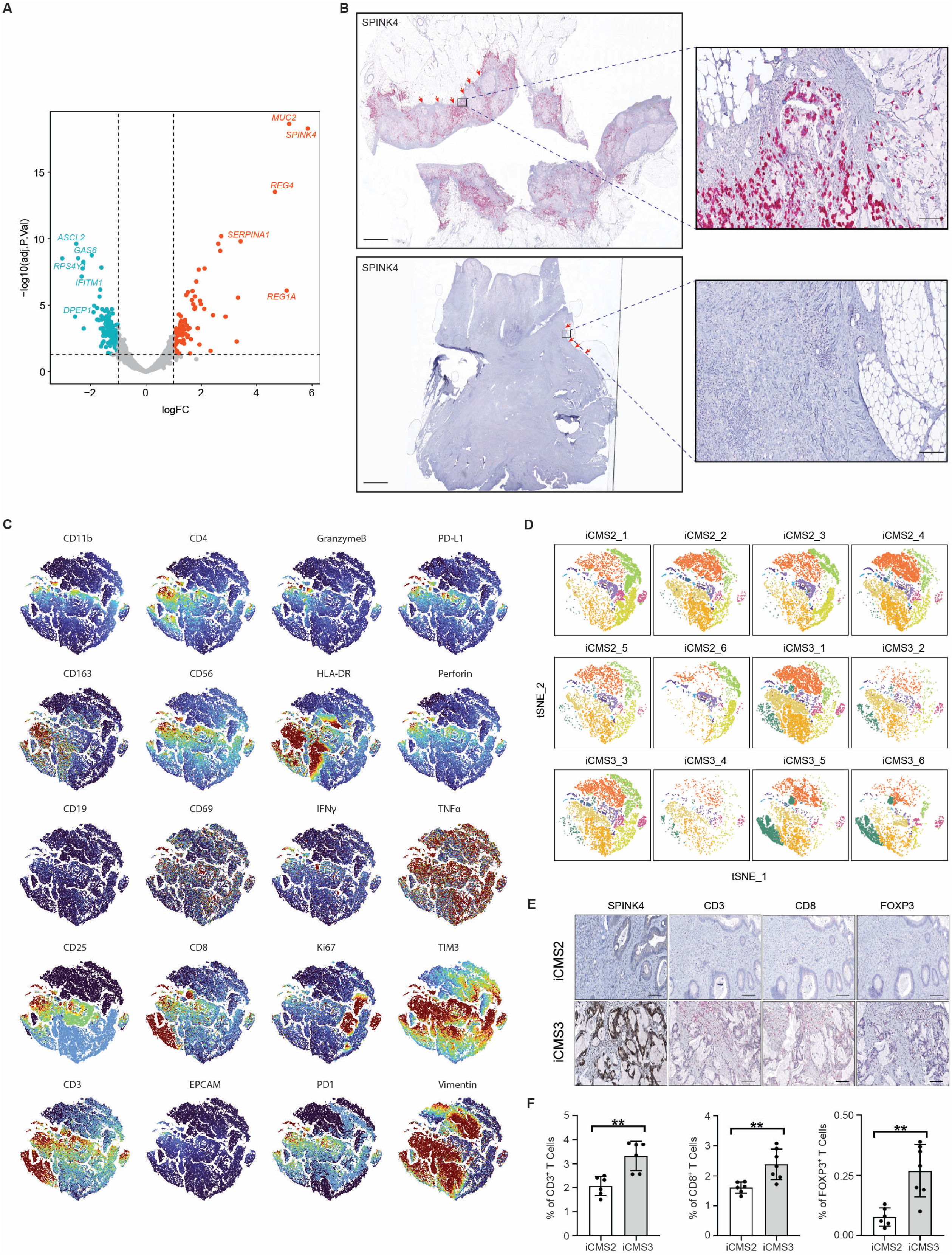
PM molecular subtypes, Related to Figure 3. (A) Volcano plot analysis showing top differentially expressed genes of the iCMS2 (blue) and iCMS3 (red) cancer cells in PM. (B) IHC staining for SPINK4 in SPINK4^+^ and SPINK4^-^ T4 primary colon cancer (red arrow, invasion border). Scale bar, 500 μm. Scale bars inside the magnification boxes, 25μm. (C) Feature plots of 20 immune markers from the imaging mass cytometry assay. (D) Imaging mass cytometry was performed in iCMS2 and iCMS3 PM samples (n=3/group, 2 images/sample) on the Hyperion platform with a total of 20 immune markers. Imaging information from all 6 samples was integrated and tSNE analysis is conducted and split into tSNE plots by each sample. (E) IHC staining for SPINK4, CD3, CD8 and FOXP3 in a cohort of PM samples (n=13). Scale bars, 50 μm. (F) Quantification of IHC staining for CD3, CD8 and FOXP3 in iCMS2 and iCMS3 PM.

**Figure S3.**
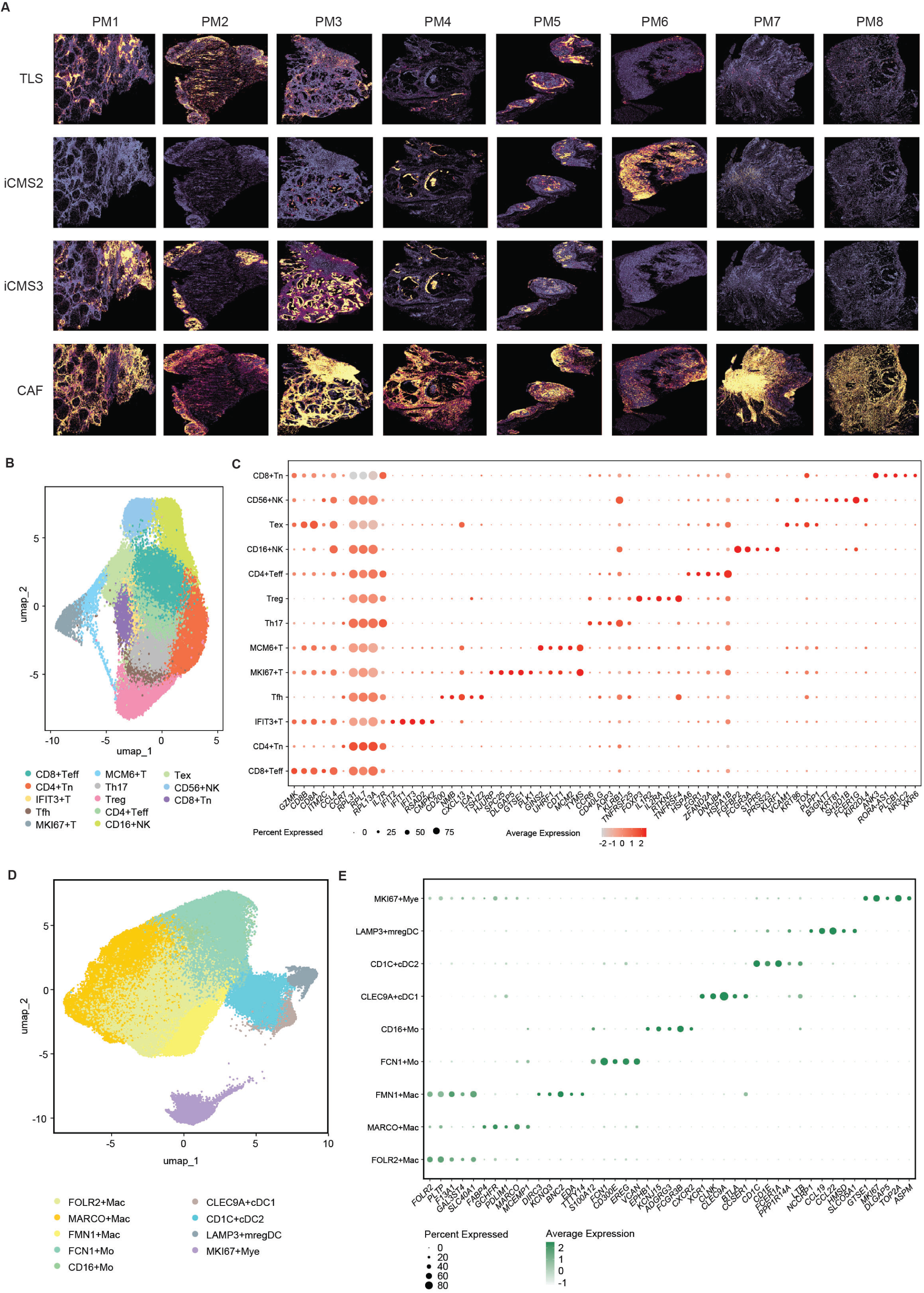
Spatial characteristics of PM, Related to Figure 4. (A) Gene signature distribution for TLS, iCMS2, iCMS3 and CAF across the 8 samples (PM1-PM8) from Xenium assays. (B) T and NK cell clusters identified from the pan-cancer scRNA-seq atlas. Tn, naïve T cell; Tex, exhausted T cell; Teff, effector T cell; Treg, regulatory T cell; Th17, T helper 17; Tfh, T follicular helper cell. (C) Maker genes for the T and NK cell clusters. (D) Myeloid cell clusters identified from the pan-cancer scRNA-seq atlas. Mye, myeloid cell; mregDC, mature regulatory dendritic cell; cDC, conventional dendritic cell; Mo, monocyte; Mac, macrophage. (E) Maker genes for the myeloid cell clusters.

**Figure S4.**
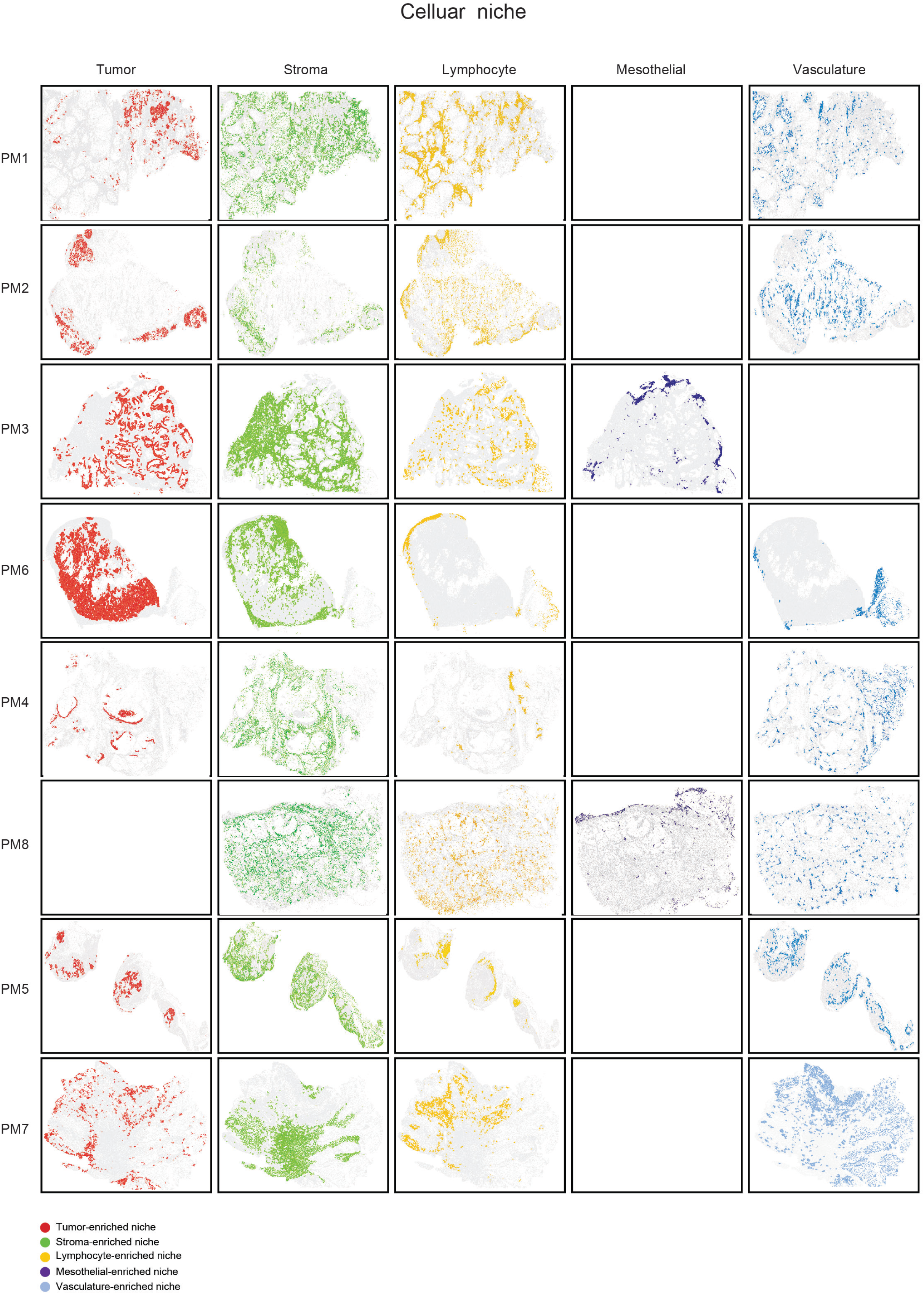
Cellular niches identified in PM samples, Related to Figure 5. Cellular niches including tumor-, stroma- and lymphocyte-enriched niches are identified in a majority of samples, while mesothelial- and vasculature-enriched niches are more context dependent.

**Figure S5.**
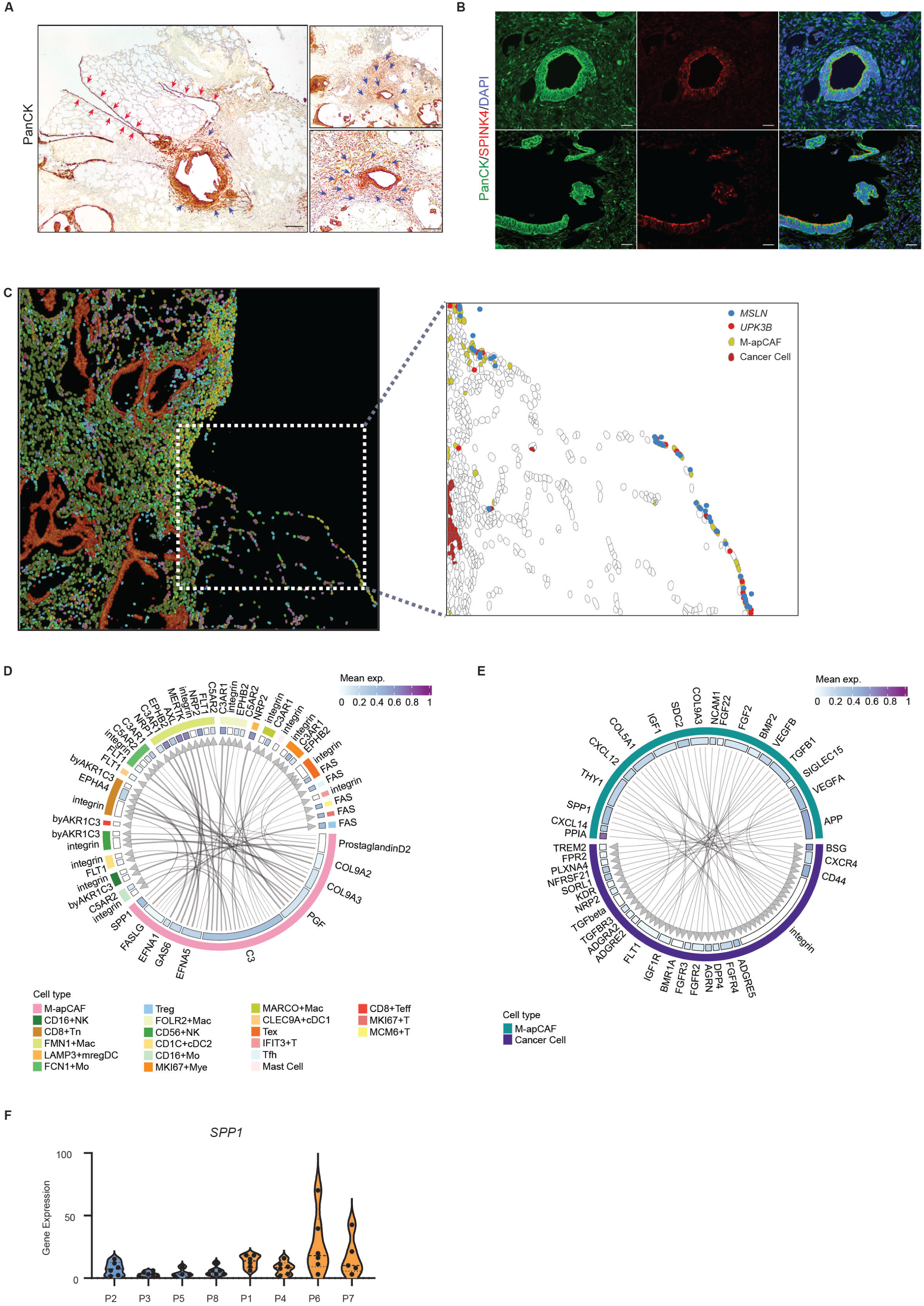
Spatial distribution and ligand-receptor interaction of M-apCAFs, Related to Figure 5. (A) IHC staining for pan-cytokeratin (PanCK) in human PM samples. Scale bars, 250 μm. (red arrow, normal mesothelium; blue arrow, cytokeratin^+^ CAFs). (B) Multiplex IHC staining for PanCK, SPINK4 and DAPI in human PM samples. Scale bars, 10 μm. (C) Visualization of normal mesothelium adjacent to M-apCAF-enriched areas from Xenium assay. M-apCAFs, cancer cells and the expression of normal mesothelial cell genes *MSLN* and *UPK3B* are shown. (D) Ligand-receptor interaction analysis between M-apCAFs (ligands) and different populations of immune cells (receptors). (E) Ligand-receptor interaction analysis between M-apCAFs (ligands) and cancer cells (receptors). (F) *SPP1* expression in the iCMS2 CAFs (patient 2, 3, 5, 8 (P2, P3, P5, P8)) and iCMS3 CAFs (patient 1, 4, 6, 7 (P1, P4, P6, P7)) from the GeoMx assay.

**Figure S6.**
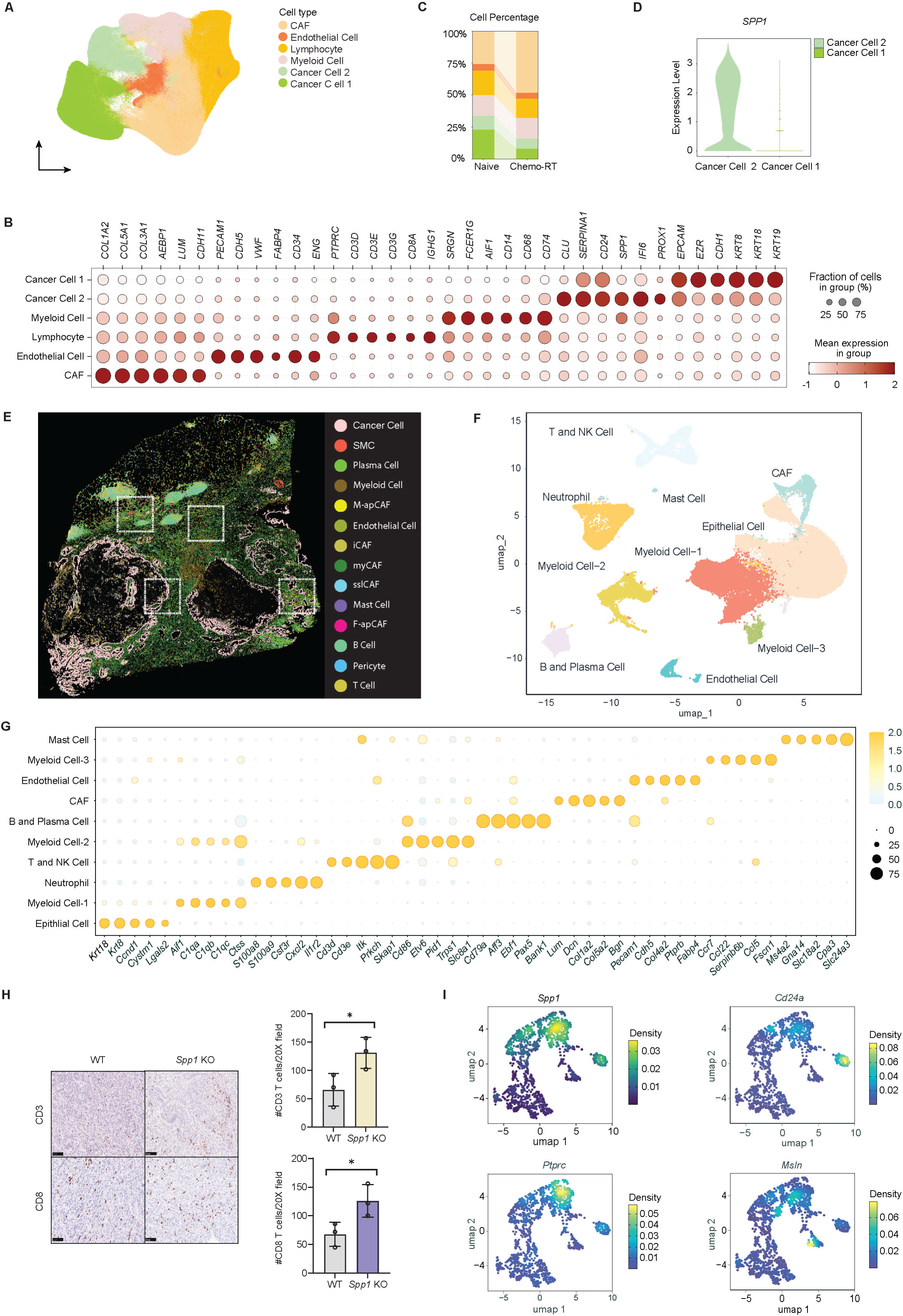
SPP1+ niches in PDAC, Related to Figure 6. (A) UMAP of major cell types across the 6 human PDAC samples identified from the Xenium assays. (B) Marker genes of the major cell types in PDAC. (C) Proportions of the major cell types in the treatment naïve and chemoradiotherapy (chemo-RT)-treated PDAC samples. (D) Expression of *SPP1* in cancer cell 1 and cancer cell 2 populations. (E) Robust cell type decomposition is performed in PDAC to deconvolve the Xenium data into cell types using our pan-cancer scRNA-seq atlas as reference. (F) *Spp1* WT or KO tumors are digested into single cell suspension and subjected to scRNA-seq. Major cell types are identified in the merged data from *Spp1* WT and KO groups. (G) Marker genes of major cell types in *Spp1* WT and KO tumors. (H) IHC staining and quantification for CD3 and CD8 in *Spp1* WT and KO tumors (n=3/group). (I) Expression of *Spp1*, *Ptprc*, *Cd24a* and *Msln* in CAFs of *Spp1* WT and KO tumors.

## Notes

### Competing Interest Statement

The authors have declared no competing interest.

